# Nitrate-responsive Mycobacterial Intracytoplasmic Membranes dampen Inflammation during *Mycobacterium tuberculosis* Infection

**DOI:** 10.1101/2025.09.17.676823

**Authors:** Camille Keck, Stéphane Tachon, Fadel Sayes, Daniele Musiani, Quentin Giai Gianetto, Mariette Matondo, Matthijn Vos, Roland Brosch, Jost Enninga

## Abstract

Subcellular compartmentalization of metabolic processes is a main feature of prokaryotic and eukaryotic architecture. Environmental bacteria generate intracytoplasmic membranes (ICMs) as a crucial strategy to adapt their metabolism to environmental changes. While pathogenic intracellular bacteria also perceive various stressful stimuli during host interactions, the subsequent re-organization of their internal architecture has not been explored. Using cryo-electron tomography, we show that *Mycobacterium tuberculosis* (Mtb), a major human pathogen, is able to form ICMs outside and inside host cells in a strain-dependent manner. We characterize these Mycobacterial intracytoplasmic Membranes (MIMs) as nitrate-induced structures involved in regulation of metabolism. Furthermore, we uncover that MIM formation during macrophage infection correlates with the ability of Mtb to dampen cellular inflammatory responses. Our findings reveal a previously uncharacterized cytoplasmic structure in Mtb and link it to a functional mechanism that enables the bacterium to adapt to its intracellular niche.

## Introduction

Historically, bacteria were thought to lack subcellular organization. Over the last years, this paradigm has shifted and it is now widely accepted that prokaryotes, like eukaryotes, compartmentalize biochemical processes via a highly sophisticated intracellular architecture. Well-known examples include bacterial microcompartments (BMCs), protein-based organelle-like structures widely distributed among Bacteria^1^. Beyond BMCs, the discovery of intrabacterial lipid-bound compartments, also named intracytoplasmic membranes (ICMs), in an increasing number of bacterial phyla highlights the wide distribution of bacterial compartmentalization^2,3,4^. The exploration of these compartments has been accelerated by novel electron microscopy methods, particularly cryo-electron tomography (cryo-ET) that enable the visualization of intracellular structures at nanometer-scale resolution^5,6^. Several bacterial phyla, especially Proteobacteria, Cyanobacteria and Verrucomicrobia, have been shown to form ICMs to spatially organize metabolic processes in response to a variety of stimuli. For example, cyanobacteria dynamically rearrange their thylakoid membrane surfaces to optimize photosynthesis efficiency under varying light conditions^7^. Similarly, purple sulfur bacteria, methanotrophs and nitrifying bacteria form ICMs to contain the components for photosynthesis, methane oxidation and nitrogen assimilation pathways ^8,9,10^, respectively. The dynamic remodeling of these structures highlights the sophistication of these species to adapt and sustain energy production in changing environments. So far, research on ICMs has focused on environmental species, however for bacterial pathogens the impact of environmental constraints during the host-pathogen crosstalk has primarily been studied at the proteome or transcriptome level^11–15^.

The intracellular bacterial pathogen *Mycobacterium tuberculosis* (Mtb), a human-adapted member of the Mtb complex (MTBC) is the causative agent of tuberculosis (TB), a disease that affected an estimated 10.8 million people worldwide in 2023 and causes over 1 million deaths each year^16^. Mtb is able to survive a variety of stressful and dynamically changing microenvironments throughout the course of infection. Macrophages represent the primary target cells of Mtb. As professional phagocytes, they internalize invading bacteria within phagosomes, a specific microenvironment that rapidly acidifies around the ingested particles^17^. Phagosomes commonly apply specific enzymatic activities, generating a superoxide burst^18^, efficiently killing many pathogens. However, studies on the transcriptional response of Mtb inside host phagosomes^11,19^ revealed a genome-wide remodeling of Mtb’s metabolism to resist and adapt to the macrophage’s hostile environment. The long history of co-evolution between pathogenic tubercle bacilli and their mammalian hosts^20^ has allowed Mtb and other MTBC members to develop an array of mechanisms to resist phagosomal stresses, blocking phagosome maturation, and promoting phagosomal rupture^21,22^ , thereby limiting the full employment of the macrophage’s hostile conditions during the infection process.

Within phagosomes, Mtb is exposed to low oxygen tension (<1%)^23^. Hypoxic conditions are further amplified by the host cell inflammatory response, which leads to the production of reactive oxygen species (ROS) and reactive nitrogen species (RNS), such as nitric oxide (NO)^24,25^. These molecules increase oxidative stress for the bacilli, exacerbating oxygen depletion, both intracellularly and extracellularly. This leads to the recruitment of neighboring immune cells to the infection site, thereby further limiting oxygen diffusion^26,27,28^. Previous transcriptomics studies have shown that such environmental signals trigger a hypoxic gene expression response in Mtb ^29^ and induce a switch from aerobic respiration to anaerobic nitrate respiration, by using nitrate as an alternative terminal electron acceptor^30^. Nitrate is readily available in hypoxic environments and has a high redox potential suitable for energy generation^31^. Most mycobacteria possess a functioning nitrate reductase (NR), and it is Mtb that exhibits the highest nitrate-reducing capacities, hinting towards a particular importance of nitrate respiration for Mtb^32^. The regulatory circuits of this metabolic adaptation have remained obscure. Anaerobic nitrate respiration has been implicated in the fitness of Mtb and *M. bovis* BCG within host cells and granulomas, although it was shown to be non-essential, as mutants lacking nitrate reduction capacity are not lethal^33,34^. However, whether basal nitrate metabolism occurs under normoxic conditions is not fully understood yet. Interestingly, nitrate *per se* was previously shown not to be a signal leading to increased expression of NR in the presence of oxygen, but low levels of NR activity were constitutively measured, indicating basal nitrate reduction even in presence of oxygen^32^. Furthermore, Mtb possesses a two-component system (TCS), NarSL (Rv0845; Rv0844c)^35^ thought to be involved in nitrate sensing and possibly nitrate metabolism regulation, although a deeper understanding of its function is lacking^36^.

Taking advantage of in situ cryo ET characterization of clinically relevant Mtb strains, we identify ICMs in this pathogen, and we propose a regulatory role of these structures in metabolic processes and adaptation to nitrate-rich environments during intracellular infection that impact the regulation of inflammatory responses within host cells.

## Results

### Differential formation of intracytoplasmic membranes (ICMs) among pathogenic mycobacteria

The relationship between morphological changes of intracellular bacterial architecture and metabolic adaptation to the surrounding environment has been characterized in many non-pathogenic bacteria^7,10^. To investigate whether a similar link between cell morphology and function exists in Mtb and other pathogenic mycobacteria, we explored differences in cellular architecture of chosen mycobacterial species. For this purpose, we performed cryo-ET on frozen-hydrated cells, coupled with cryo-FIB/SEM to thin the mycobacterial cell layer prior to cryo-TEM imaging providing windows for the observation of the cytosolic structure at the nanoscale (Fig. 1A). For our experiments, we included the widely used laboratory-adapted Mtb strain H37Rv along with a selected virulence-attenuated H37Rv ESX-1 deletion mutant^37^, as well as the highly immunogenic Mtb CDC1551, Mtb Erdman, the *Mycobacterium bovis* BCG vaccine strain, and the distantly related nonpathogenic non-MTBC member *M. smegmatis* MC^2^155. The uniform rod-shaped morphology of the bacilli in our tomograms along with the continuous and intact appearance of their cell wall were used as indicators of good sample preservation.

**Figure 1:**
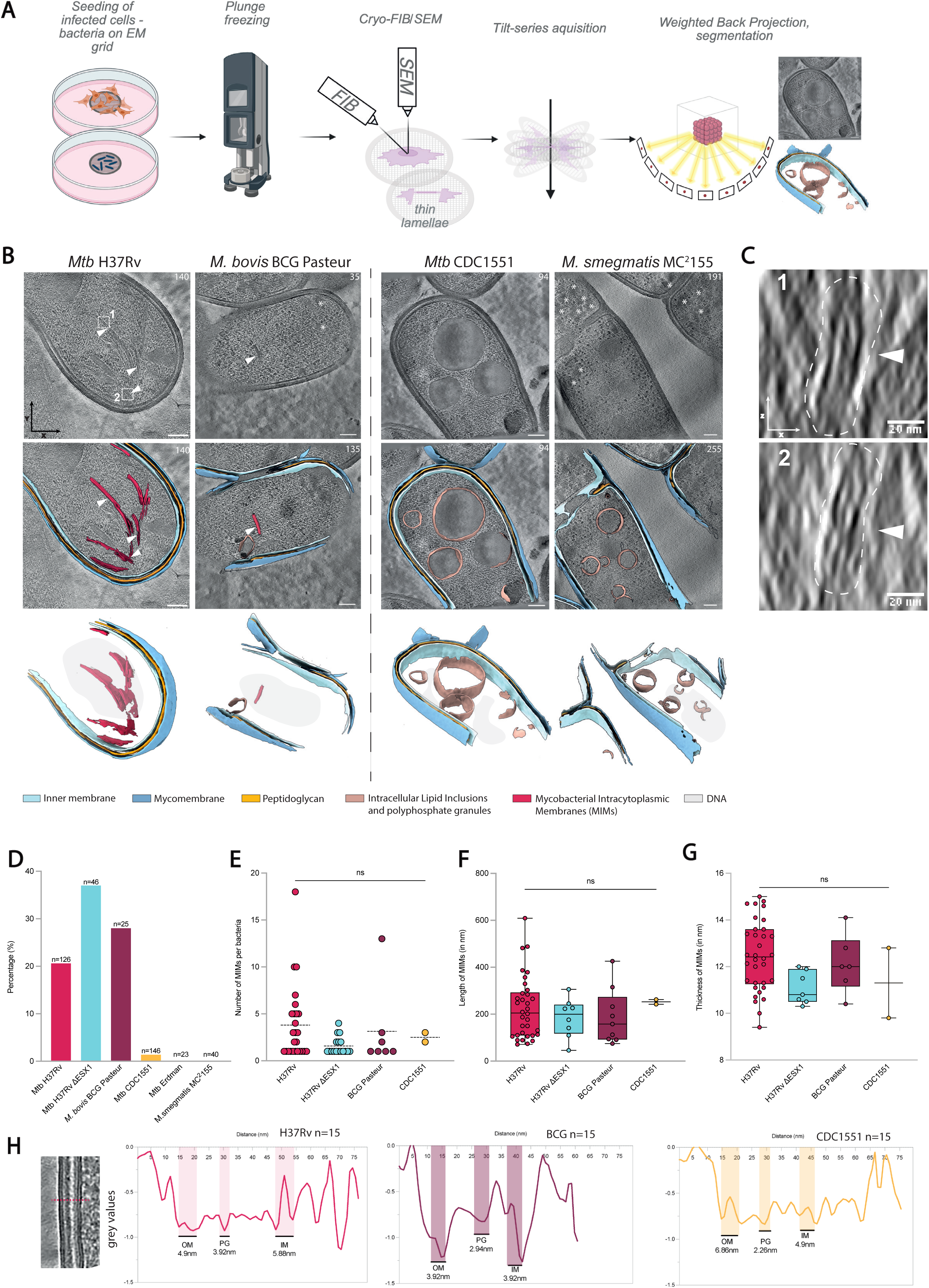
Structural organization of diverse mycobacterial species by cryo-ET and characterization of Mycobacterial Intracytoplasmic Membranes (MIMs). **A.** Graphic overview of the workflow used in this study to obtain cryo-electron tomograms of frozen-hydrated mycobacterial cells from liquid culture or from infected eukaryotic cells. Bacterial suspension or infected cells are deposited on EM grids and plunge frozen in liquid-nitrogen cooled ethane. The frozen bacterial layer or infected cells are thinned by cryo-FIB milling. Tilt-series are acquired with a transmission electron microscope, and volumes are reconstructed with a weighted back projection algorithm. **B.** Tomographic slices and their corresponding volume segmentations of Mycobacterium tuberculosis H37Rv, CDC1551, M. bovis BCG Pasteur and M. smegmatis MC^2^155, harvested from exponential phase liquid cultures. White arrowheads: MIMs. White stars: α-glucans storage granules. Tillt-series were acquired at 42 000x (Mtb H37Rv and M. smegmatis MC^2^155), or 53 000x (Mtb CDC1551 and M. bovis BCG Pasteur). Z-slices are shown in top right corner. Scale bars: 100nm. **C.** Zoom-in of insets 1 and 2 of figure 1B (Mtb H37Rv) oriented in the XZ axis. Scale bar: 20nm. **D.** Quantification of MIMs in mycobacterial cells imaged by cryo-ET. Percentage represents the fraction of cells of each strain in which MIMs were visible. **E.** Number of MIMs counted per MIM-positive bacilli. Kruskal-Wallis tests with Dunn’s multiple comparisons tests were performed. ns: non-significant. **F.** Length of MIMs in nm, measured in IMOD. Kruskal-Wallis tests with Dunn’s multiple comparisons tests were performed. ns: non-significant. G. Thickness of MIMs in nm, measured in IMOD. Kruskal-Wallis tests with Dunn’s multiple comparisons tests were performed. ns: non-significant. **H.** Density profiles of mycobacterial cell wall in corresponding species. The profiles were calculated by averaging cross-sections of the cell envelopes in 15 different bacilli for each species. OM: outer membrane; PG: peptidoglycan; IM: inner membrane. Z-slices number are shown on the top left corner. Scale bar: 50nm.

Our cryo-tomograms revealed a diverse array of previously described features of mycobacterial cells, including the waxy lipid-rich characteristic mycomembrane, the peptidoglycan layer and the phospholipid bilayer of the inner membrane (IM) (Fig. 1B, Supplementary Movie 1)^38,39^. We also observed in all above-mentioned mycobacterial strains intracellular lipid inclusions (ILI), polyphosphate granules (PP), α-glucan storage granules (Fig. 1B, Supplementary Fig. 1), and the ribosome-exclusion zone around the nucleoid (Supplementary Fig. 1, Supplementary Movie 2), consistent with previous observations^40^.

Importantly, our cryo-tomograms revealed the presence of unidentified membrane-bound compartments, observed exclusively within the cytosol of the investigated pathogenic mycobacteria (Fig. 1B, Supplementary Fig. 2). These cytoplasmic structures appeared as long and thin membranes when viewed on the XY axis (Fig. 1B). For both Mtb and for *M. bovis* BCG, they exhibited homogeneous thickness in XY and XZ (Fig. 1C) indicating flattened sheet-like structures oriented parallel to the cell’s length (Supplementary Fig. 3A). Comparing our newly identified intracellular structures with available electron micrographs from other bacteria, they represented all features of ICMs, a cytosolic bacterial compartment that has been described for various environmental bacteria. Therefore, we decided to name these structures as Mycobacterial intracytoplasmic Membranes (MIMs).

The MIMs appear as long sheets mostly along the parallel axis of the bacilli, although not exclusively (Supplementary Fig. 3B). ICM sub-cellular localization in environmental bacteria has been shown to be species-specific, with some purple non-sulfur bacteria species developing polar ICMs^8^, similarly to recently observed *Coxiella burnetti* ICMs^41^, and photosynthetic cyanobacteria developing thylakoids all around the periphery of the cell^7^. In our tomograms, we found that approximately 60% of MIMs were localized throughout the cell body while the remaining 40% were localized close to the cell pole (Supplementary Fig. 3C). We also noted that in most of our tomograms, all MIMs within a given bacillus appeared as grouped bundle close to one another, rather than being scattered throughout the cytoplasm. Finally, some of the MIMs appeared to have a certain degree of membrane curvature, while others were almost entirely straight (Fig. 1B).

We quantified the amount of MIMs in all analyzed strains. This revealed their presence in 20.6% of all *M. tuberculosis* H37Rv bacilli imaged by cryo-ET (Fig. 1D). Notably, MIMs were also observed in *M. tuberculosis* H37Rv ΔESX-1 and *M. bovis* BCG Pasteur, both lacking the ESX-1 type VII secretion system (T7SS)^42^, indicating that the formation of MIMs is independent of ESX-1 functions.

A major difference in the presence of MIMs was found for Mtb CDC1551 (only 1.4% exhibited MIMs) and their absence in Mtb Erdman, although we cannot rule out their presence in the latter due to the relatively low number of tomograms of Mtb Erdman acquired (Fig. 1D). It should be noted that Mtb H37Rv, CDC1551 and Erdman all belong to the modern Euro-American lineage 4 of the MTBC. While these strains exhibit very high levels of genomic sequence identity^43^, they diverge in their pathogenicity and immunogenicity, with CDC1551 being reported to be highly immunogenic ^44,45,46^. Finally, no MIMs were found in the non-pathogenic non-MTBC strain *M. smegmatis* MC²155, suggesting that MIMs might be a unique feature of MTBC members, showing differential degrees of adaptation to the host cell environment.

The number of MIMs per bacterium ranged from 1 to 18, though this likely underestimates their exact numbers, as our reconstructed tomograms covered only about one-fifth of the bacterium’s volume, and the magnification used did not allow full visualization of each cell’s full length (Fig.1E). While most of the MIMs observed measured between 100 and 300nm in length, some were as long as 600nm, and the global length of all MIMs as well as their thickness measured in all strains did not vary significantly (Fig. 1F, Fig. 1G). Furthermore, the thickness of the MIMs consistently followed the same trend across all MIM-positive strains, being roughly twice that of the inner membrane (Fig.1G, 1H) of each respective strain. This consistent ratio points towards the possibility that MIMs are derived from the IM phospholipid bilayer rather than being formed *de novo*.

Altogether, our cryo-tomogram analysis shows that mycobacteria form intracytoplasmic membranes differentially between strains with a morphology that is conserved across MIM-positive strains. Their exclusive presence in the investigated pathogenic strains suggests a role during host cell infection.

### MIM formation is a dynamic process in response to environmental stimuli

To investigate whether MIM formation is differentially regulated during infection, we acquired tomograms of intracellular Mtb following infection of RAW264.7 macrophages with both H37Rv and CDC1551 (Fig. 2A). We found that the proportion of intracellular MIM-positive H37Rv decreased 2.4-fold compared to its extracellular counterpart (Fig. 2B). In contrast, the proportion of MIM-positive CDC1551 cells increased 4.7-fold in the intracellular environment relative to extracellular CDC1551 (Fig. 2B). Furthermore, using cryo-CLEM allowed us to target infected cells specifically during or after phagosomal rupture. Consequently, we were able to acquire cryo-tomograms of both phagosomal and cytosolic Mtb, allowing us to capture a broader range of stress conditions associated with each intracellular niche and to assess their impact on MIM formation (Supplementary Fig. 4A). We found that MIM-positive H37Rv were exclusively cytosolic, whereas MIM-positive CDC1551 were present in both the phagosome and the cytosol of infected macrophages (Supplementary Fig. 4B). Finally, MIM sizes between *in vitro* culture and intracellular condition did not vary significantly (Fig. 2C), indicating that while the formation of MIMs was sensitive to the specific environmental context, their morphology was independent of it.

**Figure 2:**
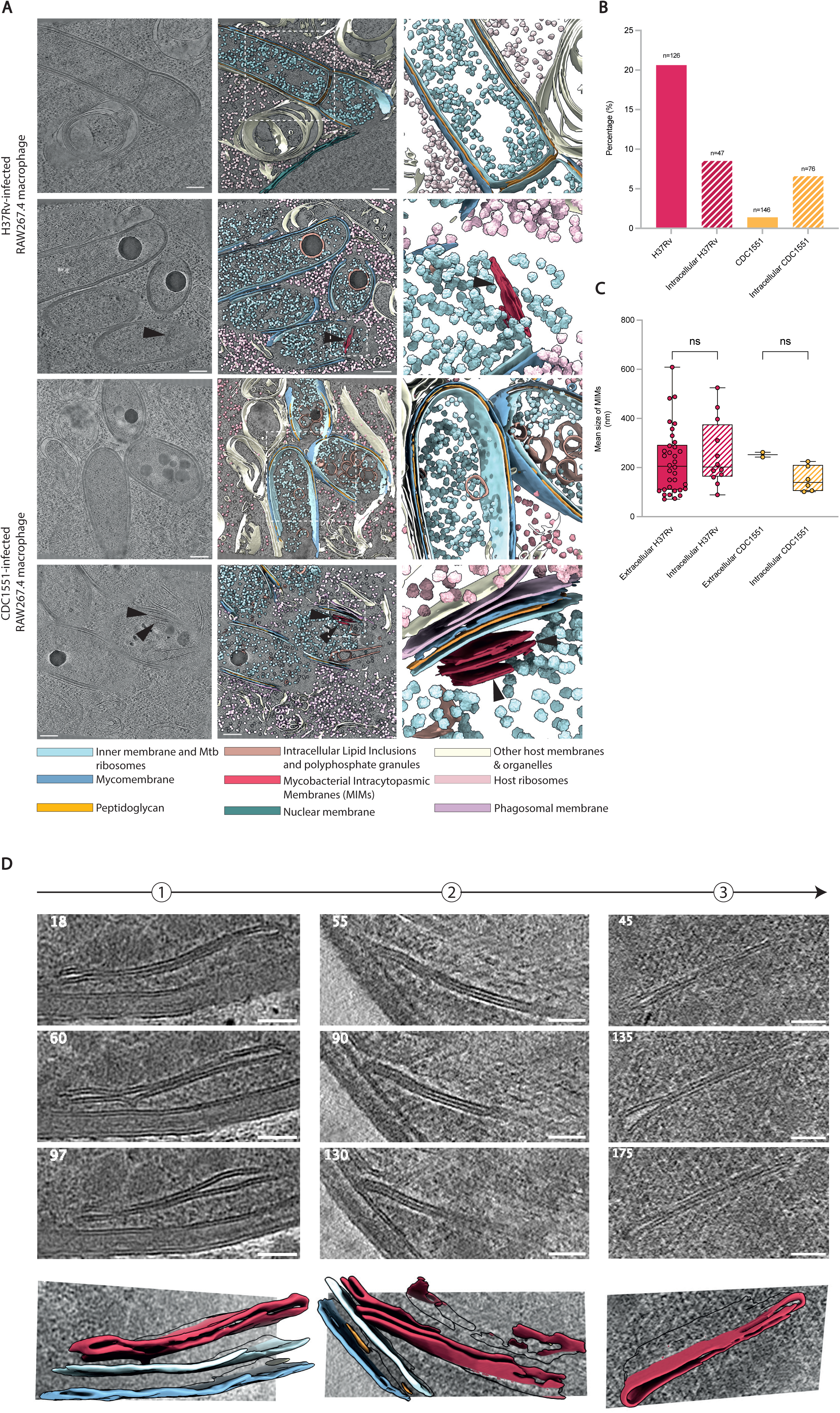
Dynamics of MIM formation. **A**. Tomographic slices of Mtb H37Rv or Mtb CDC1551-infected RAW264.7 macrophages. Segmentations are shown for each inset. Black arrowheads show mycobacterial MIMs. Scale bars: 200 nm. **B**. Quantifications of intracellular MIM-positive H37Rv and CDC1551 in comparison with MIM quantifications of independently, previously acquired tomograms of in vitro Mtb, as shown in Fig. 1. **C**. Length of MIMs found in intracellular Mtb in comparison with MIM length in independently, previously acquired tomograms of in vitro Mtb, as shown in Fig. 1. Cryo-tomograms were acquired in at least two independent experiments. Mean, SD and individual values are shown. Ordinary Two-Way ANOVA with Tukey’s multiple comparisons test was performed. ns: non-significant. **D**. Montage of tomographic slices representing different stages of MIM formation and their corresponding segmentations. Stars show contact point between MIM and IM. Z-slices number are shown on the top left corner. Scale bar: 50nm.

### These results suggest that the basal turnover of MIMs observed *in vitro* is exacerbated during intracellular infection

To investigate MIM biogenesis, we analyzed their morphology and their interaction with the surroundings in more detail by cryo-tomography, especially with regards to the IM. To this end, we acquired a total of 374 tomograms (all strains included) of liquid-culture grown cells harvested in exponential phase. This allowed an in-depth characterization of MIM formation and architecture from 651 individual bacilli. Of these, 138 were MIM-positive. The reconstructed volumes captured different stages of MIM formation. While most appeared as freely floating structures in the cytosol, distant from the inner membrane, some displayed clear continuity with the IM, exhibiting a sheet-like compartment that remains connected to the IM (Fig. 2D, Supplementary Fig. 4C). Other MIMs were observed in close proximity to the IM, with their tips either forming near-contact points or lacking visible connections yet still positioned close to the IM (Fig. 2D, Supplementary Fig. 4C). Although cryo-ET captures static images representing structures at a single time point and thus lacking temporal resolution, our observations support a stepwise process for MIM formation with a gradual invagination of the phospholipid bilayer of the IM until full detachment.

Together, these findings suggest that MIMs are dynamically formed and may be actively recruited to the IM during infection in response to host-derived signals, or their formation is modulated depending on the intensity of the signal received, highlighting the dynamic turnover of these structures in response to environmental cues. Moreover, the distinct MIM levels observed in H37Rv and CDC1551, both in culture and during host infection, indicate that they reacted with different amplitudes hinting at differing sensitivities or regulatory thresholds.

### Identification of nitrate metabolism-related proteins within MIMs

Bacterial ICMs foster diverse protein complexes that support responses to specific signals. We hypothesized that MIMs in Mtb may contain proteins responsive to environmental cues. To analyze the proteome of the MIM-enriched fraction, we fragmented H37Rv and CDC1551 by mechanical lysis and density gradient ultracentrifugation (Fig. 3A) as described previously⁴⁴, isolating the MIM-rich fraction from H37Rv and the near MIM-free equivalent fraction from CDC1551. As MIMs are extended cytosolic membranes, we expected them in the denser cytoplasmic fraction of the gradient. We confirmed their presence in our enriched fraction by negative-stain TEM (Supplementary Fig. 5). The proteome from this fraction of H37Rv and CDC1551, as well as the equivalent fraction from *M. smegmatis* MC²155 as MIM-less negative control, were analyzed by mass spectrometry. Through this, we identified 2898 proteins across the five replicates of H37Rv and CDC1551 (Fig. 3B). Consistent with the high genetic similarity between H37Rv and CDC1551, only 10 proteins were exclusively found in H37Rv, and 11 in CDC1551 (Supplementary Table 1A and 1B). We then filtered the list of proteins to only keep proteins present in ≥4 of 5 replicates and supported by >2 unique peptides in ≥n–1 replicates, which yielded a list of 938 proteins (Supplementary Figure 6A). The gene ontology (GO) term analysis for molecular functions of the filtered protein list revealed an enrichment of components associated with O-methyltransferase activity, aldehyde dehydrogenase (NAD+) activity, quinone binding, acyl carrier protein synthase activity and oxidoreductase activity, corresponding to activities usually related to energy metabolism (Fig. 3C). 35 candidates were significantly enriched in H37Rv (Supplementary Figure 6A, Fig. 3D, Supplementary Table 2) with a fold-change > 2, and 20 in CDC1551 (Supplementary Figure 6A, Supplementary Table 3), with a fold-change <-2. To identify putative proteins associated with MIMs, we examined H37Rv-enriched proteins with predicted transmembrane domains in DeepTMHMM v1.0.0^47^. Only two were identified: Rv0072, involved in glutamine import^48^, and NarS (Rv0845c), the sensor kinase of the NarS/NarL two-component system (TCS)^36^. NarS being more enriched in H37Rv than Rv0072 (∼4.7 fold and ∼2.69 fold, respectively; Supplementary Table 2), NarS was selected for further analysis. We quantified mRNA expression of *narS*, its cognate transcriptional regulator *narL*, and other redox-associated genes from our mass spectrometry analysis by qRT-PCR and compared their relative expression between H37Rv and CDC1551 (Fig. 3E). Although *narS* expression was globally higher in H37Rv, the difference was not statistically significant due to inter-replicate variability. However, *narL* expression was significantly, albeit modestly, more elevated in H37Rv (Fig. 3E), although our proteomics analysis did not show NarL as significantly enriched in neither H37Rv nor CDC1551 (Supplementary Fig. 6B). None of the other redox-associated genes tested showed differential expression between the two strains at the mRNA level (Fig. 3E). These results led us to hypothesize that NarS might be associated with MIMs.

**Figure 3:**
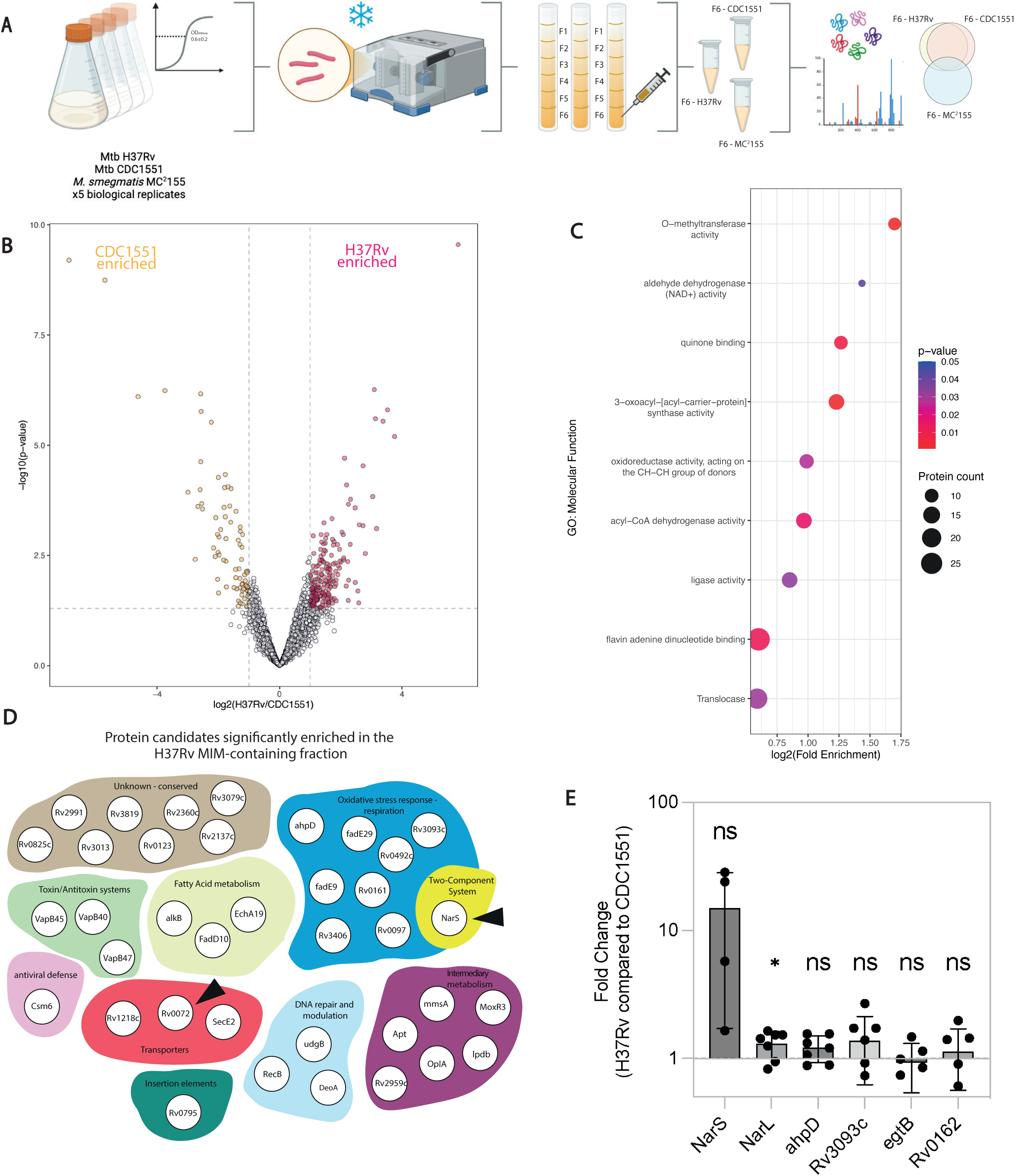
**Proteome of the MIM-enriched Mtb H37Rv fraction**. **A**. Graphic representation of the pipeline used to fragment bacterial cultures for mass spectrometry analysis of the MIM-containing fraction. Details are provided in the corresponding methods section. **B**. Volcano plot of proteins identified in all 5 biological replicates of the MIM-containing fraction of H37Rv and the equivalent fraction of CDC1551. Proteins significantly enriched in H37Rv are labeled in pink. Proteins significantly enriched in CDC1551 are shown in yellow. Non significantly enriched proteins are shown in grey. **C**. Dot plot depiction of GO analysis for molecular functions using all proteins found in 4/5 replicates with more than 2 unique peptides in n-1 replicates. Only significant terms are shown. Protein count represents the number of proteins in the GO term. Analysis was performed with the functional clustering tool on DAVID with medium classification stringency. **D**. Cloud of protein candidates significantly enriched in the H37Rv MIM-containing fraction compared to equivalent fraction in CDC1551. Proteins are clustered in categories representative of their functions. Black arrowheads show transmembrane proteins identified by DeepTMHMM. **E**. Expression of redox-related genes by qRT-PCR. Gene expression is compared as fold change between H37Rv and CDC1551, both normalized to rpoB expression in their own genetic background. Experiment was performed on 2 biological replicates. Mean and SD are shown. One sample t-tests were used with comparison of the fold change to 1. * indicates p value < 0.05, ** <0.01, *** <0.001 and **** <0.0001. ns : non-significant.

Although previous research on NarS/NarL in mycobacteria is scarce, its sequence identity to the homologous nitrate-sensing TCS NarQ/NarX and NarX/NarL in *E. coli* ^49^ and *P. aeruginosa*^50^ respectively support the role of NarS/NarL in nitrate metabolism. It is of note that response regulators of paired TCS usually bind to the promoter region of their own gene as well as that of the sensor histidine kinase. However, NarL was previously shown to lack a binding site in the intergenic region between *narS* and *narL* (Supplementary Fig. 6C), a finding supported by AlphaFold3 structural predictions of *narL* binding to the 84bp nucleotide region between *narS* and *narL* (Supplementary Fig. 6D). In contrast, co-interaction models of NarL with DosR, the response regulator of the DosRST TCS, with which NarL has previously been shown to interact^36^, predict a putative binding interface within this intergenic region (Supplementary Fig. 6D), suggesting a potential co-regulation of the NarSL TCS with DosRST.

Together, our results show that a large proportion of the proteins identified in the MIM-containing fraction are involved in redox processes, including redox sensing and oxidoreductase activity. Notably, NarS is significantly more abundant in the MIM-containing fraction of H37Rv compared to the equivalent fraction in CDC1551, which we have shown lacks MIMs when grown *in vitro*.

### Ambient nitrate regulates MIM formation

To investigate the potential link between MIM formation and nitrate metabolism in Mtb, we cultured Mtb H37Rv and CDC1551 under conditions with gradually increasing concentrations of nitrate (0, 10, 50 and 100mM). We found that both strains exhibited similar growth patterns until mid-logarithmic phase, at which point CDC1551 started showing a growth defect when cultured in 10mM NaNO_3_, the lowest nitrate concentration used, while this concentration had no impact on H37Rv growth compared to the control condition at the same timepoint (Fig. 4A). This was reflected by the significantly lower area under the growth curve (Fig. 4B) for CDC1551 at all nitrate concentrations used compared to the control condition without nitrate. This was not observed in H37Rv at any of the nitrate concentrations used. The colony forming units (CFU) counts further supported this trend with no visible impact of nitrate addition until day 4 in either strain, and a decrease in CFU counts in CDC1551 from day 7 onward when cultured with the lowest nitrate concentration that appeared further accentuated at higher concentrations (Fig. 4C). In contrast, in H37Rv we observed an increased CFU count at high nitrate concentrations, suggesting that Mtb might use nitrate as an alternative energy source when nutrients become depleted, particularly at later timepoints during growth (Fig. 4C). While these differences were consistently observed, they did not reach statistical significance due to high inter-replicate variability. These results suggested that either H37Rv tolerates high nitrate concentrations better than CDC1551, or alternatively CDC1551 undergoes extensive cellular rearrangement when grown under elevated nitrate, leading to the observed fitness cost during growth.

**Figure 4:**
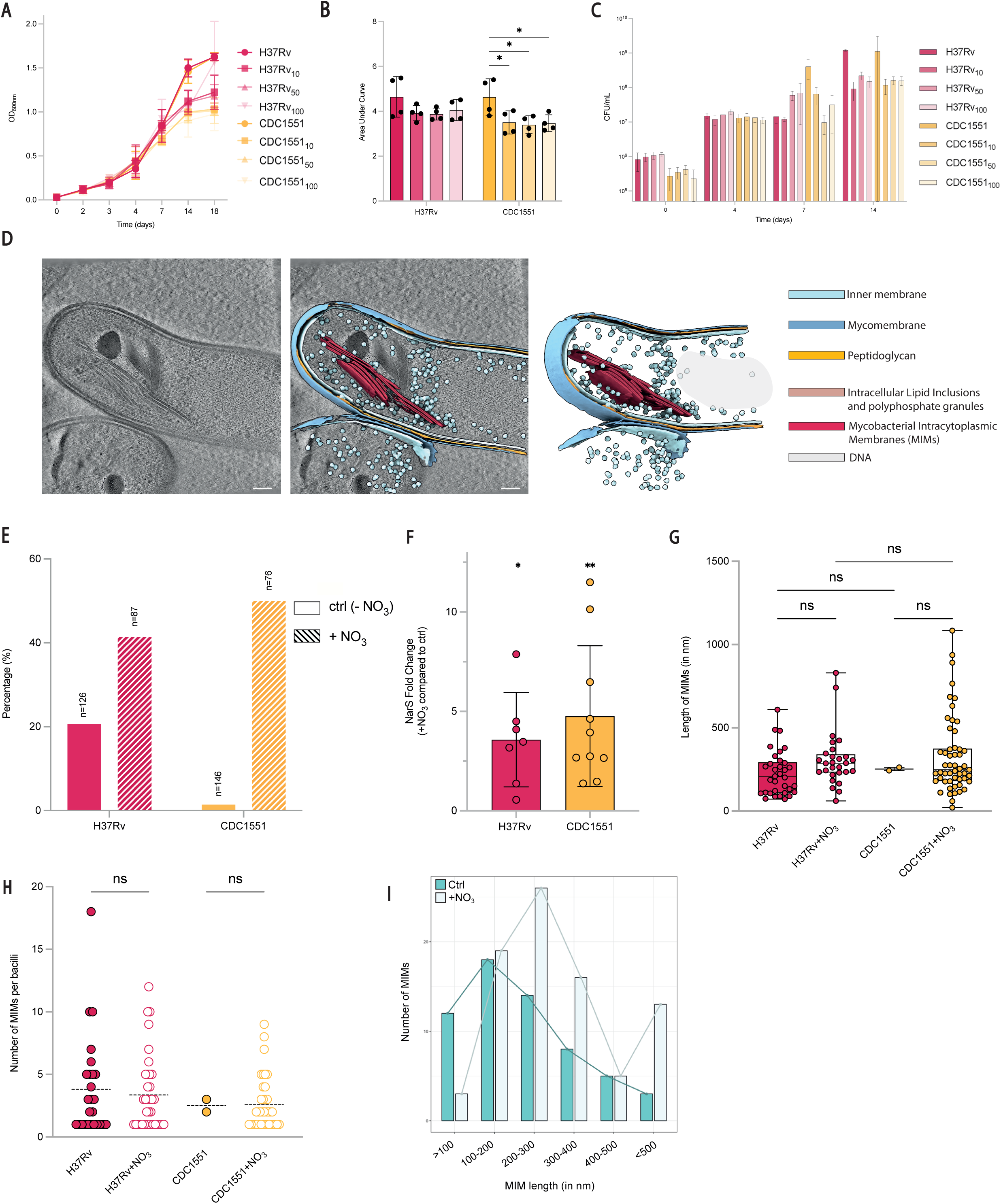
Nitrate acts as a MIM-inducing signal in Mtb H37Rv and CDC1551. **A**. Growth curves of Mtb H37Rv and CDC1551 grown in absence or presence of 10mM, 50mM or 100mM NaNO_3_. Experiment was performed on two biological replicates. Mean and SD are shown. **B**. Corresponding Area Under Curve (AUC) of growth curves shown in A. Individual values, mean and SD are shown. * indicates p value < 0.05, ** <0.01, *** <0.001 and **** <0.0001. **C**. Colony Forming Units (CFU) of corresponding samples analyzed in A. and B. Mean and SD are shown. **D**. Representative tomographic slice of Mtb cultured in 100mM NaNO_3_ for 6 days. Scale bar: 100nm. **E**. Quantification of MIMs in mycobacterial cells cultured with 100mM NaNO_3_ compared to control conditions, obtained independently in previous acquisitions as shown in Fig.1. Percentage represents the fraction of cells of each strain in which MIMs were visible. **F**. NarS expression quantified by qRT-PCR in H37Rv or CDC1551. Gene expression was normalized to the expression of rpoB in the corresponding genetic background in each condition. Comparison of gene expression is shown as fold-change between +NO_3_ condition related to the control condition without NO_3_. Experiment was performed on 2 biological replicates. One sample t-tests were used with comparison of the fold change to 1. * indicates p value < 0.05, ** <0.01. **G**. Length of MIMs in nm, measured in IMOD. Individual values are shown. Kruskal-Wallis with Dunn’s multiple comparisons tests were performed. ns: non-significant. **H**. Number of MIMs counted per MIM-positive bacilli. Means and individual values are shown. Kruskal-Wallis with Dunn’s multiple comparisons tests were performed. ns: non-significant. **I**. Distribution of MIM lengths in control growth conditions and in NO_3_ growth condition. H37Rv_10_: H37Rv, 10mM NaNO_3_. H37Rv_50_:H37Rv, 50mM NaNO_3_. H37Rv_100_: H37Rv, 100mM NaNO_3_. CDC1551_10_: CDC1551, 10mM NaNO_3_. CDC1551_50_: CDC1551, 50mM NaNO_3_. CDC1551_100_: CDC1551, 100mM NaNO_3_.

Therefore, we investigated the impact of changing nitrate concentrations in the environment of Mtb on MIM formation at the ultrastructural level. We harvested cultures of both H37Rv and CDC1551 after 6 days of growth with 100mM NaNO_3_ and imaged them using cryo-ET as described above (Fig 4D, Supplementary Fig. 7). We observed a substantial increase in MIM content in both strains, with a 2-fold increase in H37Rv, and a striking 52-fold increase in CDC1551 cells containing MIMs, compared to Mtb grown without nitrate (Fig. 4E). We hypothesized that this strong increase in MIM formation in CDC1551 might contribute to the observed growth defects, as the formation of such extensive membrane structures could be energetically costly. Importantly, addition of sodium nitrate to the medium did not affect the pH of the culture overtime (Supplementary Fig. 8A), nor compromise the integrity of the mycobacterial membrane, as shown by an Ethidium Bromide (EtBr) accumulation assay (Supplementary Fig. 8B). We thus conclude that formation of MIMs in the elevated nitrate condition was specifically induced by nitrate itself. Accordingly, expression of narS which we identified as major component of enriched MIM fraction was increased 3.5-fold in H37Rv and 4.7-fold in CDC1551 when strains were grown under 100mM NaNO_3_ (Fig. 4F), suggesting positive regulation of *NarS* in response to nitrate in the environment. Furthermore, we showed that *dosR* and to a lesser extent *dosS*, but not *dosT*, were significantly overexpressed in both H37Rv and CDC1551 in presence of 100mM NaNO_3_ in the growth medium (Supplementary Fig. 9), further strengthening the link between the expression of NarSL and MIM formation via a connection with the DosRST regulatory pathway. Although nitrate-induced MIMs were neither significantly longer within individual bacilli (Fig. 4G) nor more abundant (Fig. 4H), compared to the MIMs in control conditions, the distribution of MIM lengths indicated that nitrate exposure leads to a higher total proportion of longer MIMs (Fig. 4I).

Together, these results reveal nitrate as a signal strongly inducing the formation of MIMs in CDC1551, and to a lesser extent in H37Rv at equivalent concentration, indicating differential regulatory thresholds for nitrate-triggered MIM formation in the two strains, with CDC1551 being more sensitive to nitrate-rich environment. The addition of nitrate to the medium increases the expression of *narS* along with its response regulator *narL*, possibly through co-regulation with other transcriptional regulators, particularly the nitric oxide responsive TCS regulator DosR.

### Induction of MIMs limits inflammation during macrophage infection

During infection of host macrophages by Mtb, inflammation levels rise rapidly to alert and recruit other immune cells to the site of infection. As such, dampening the inflammatory response is considered advantageous for the pathogen. To examine the role of Mtb MIMs on infection outcomes, we infected RAW264.7 macrophages with Mtb H37Rv or CDC1551 strains grown under control conditions or in the presence of 100 mM nitrate, corresponding to basal or elevated MIM levels, respectively. We first confirmed that Mtb internalization was similar comparing both strains and conditions, indicating that MIMs do not influence phagocytosis of the bacilli (Fig. 5A). Over a 7-day period, intracellular replication resulted in increased CFU counts for both tested Mtb strains, but no significant differences were observed between the control and added nitrate conditions at any time point post-infection (Fig. 5A). We therefore concluded that MIMs do not significantly affect internalization or intracellular replication. Furthermore, we reasoned that MIMs in Mtb do not induce an increase of host-mediated bacterial killing as CFU counts continued to rise throughout the measured time courses.

**Figure 5:**
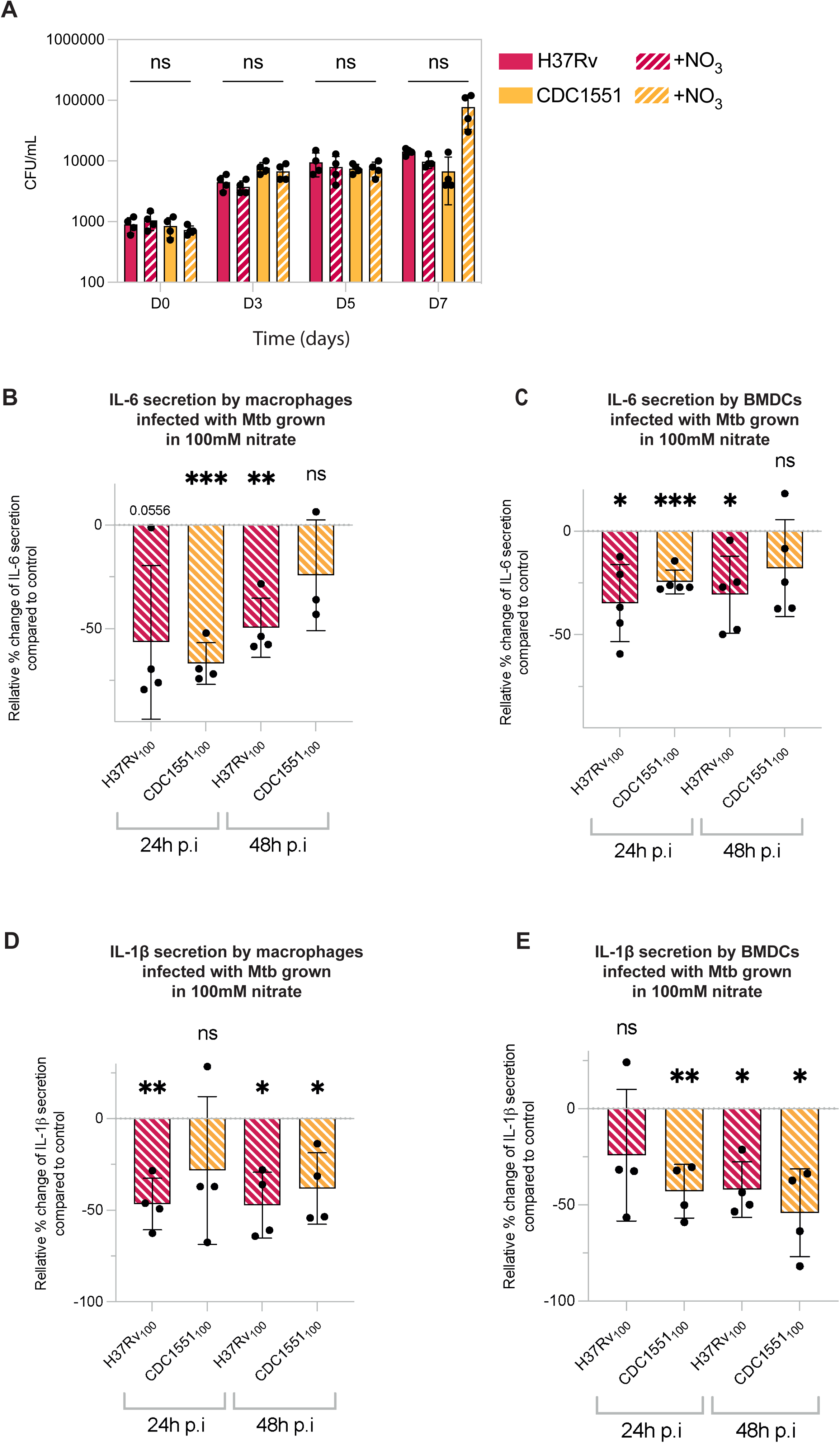
induction of MIMs in Mtb limit pro-inflammatory responses in infected cells. **A**. Colony forming units of intracellular Mtb during 7 days of RAW264.7 macrophage infection. Individual CFU/mL values and means are shown. Statistics were calculated using two-way ANOVA and Tukey’s multiple comparisons test. ns: non significative. **B.-E:** Cytokine quantifications by ELISA assay. Results are expressed as the relative percentage change in IL-6 (B, C) and IL1β (D, E) secretion by macrophages (B, D) or Bone Marrow-derived Dendritic Cells (C, E) infected with Mtb grown in the presence of 100 mM NaNO_3_, compared with Mtb grown under control conditions (without NaNO_3_). Individual values, mean and SD are shown. Experiments were performed on biological duplicates. One sample t-tests were used with comparison to 0. * indicates p value < 0.05, ** <0.01, *** <0.001. ns: non-significant

We shifted our focus towards possible effects of MIMs on the infected macrophages and their inflammatory responses, given the reported differences in immunogenicity of strains Mtb H37Rv and CDC1551^51^. To this end, we examined how MIM formation influences inflammatory responses after host cell infection with both strains measuring the release of pro-inflammatory cytokines typically induced during Mtb infection. Among these, interleukin-6 (IL-6) is particularly important for host control of infection^52^. Quantification of IL-6 secretion was consistently and significantly reduced when cells were infected with both H37Rv or CDC1551 cultured in 100 mM nitrate (MIM-induced) compared to their counterparts grown under control conditions (Fig. 5B, Fig. 5C). Specifically, infection of macrophages with MIM-induced H37Rv reduced IL-6 secretion by 56.6% at 24 hours post-infection (p.i.) and by 49.6% at 48 hours p.i, while infection with MIM-induced CDC1551 reduced IL-6 secretion by 66.8% at 24 hours p.i. and 24.2% at 48 hours p.i (Fig. 5B).

Similar trends were observed in BMDCs. Infection with MIM-induced H37Rv decreased IL-6 secretion by 34.7% at 24 hours and 30.7% at 48 hours p.i, while CDC1551 under the same conditions reduced IL-6 by 24.5% at 24 hours, with no significant effect at 48 hours p.i (Fig. 5C). These findings suggest that nitrate–induced MIM formation significantly dampens IL-6 secretion during Mtb infection. This effect is most pronounced at 24 hours post-infection and is more potent with CDC1551 than with H37Rv, reflecting the stronger MIM formation observed in CDC1551.

The overall decrease in IL-6 secretion by MIM-induced Mtb was also observed for other pro-inflammatory cytokines, for IL-1β in both cell types (Fig. 5D, Fig. 5E) and for TNFα to a lower extent, mainly in macrophages (Supplementary Fig. 10A). Additionally, we also observed a modest but significant increase of the anti-inflammatory cytokine IL-10 secretion by RAW macrophages when infected with MIM-induced H37Rv, at 48 hours p.i, and by BMDCs when infected with MIM-induced CDC1551 at 24 hours p.i.

Together, these results indicate that while nitrate-induced MIM formation in Mtb does not affect bacterial uptake or replication, the presence of MIMs correlates with the modulation of cellular immune responses highlighted by the dampening of inflammation. Notably, the decrease of pro-inflammatory responses was particularly pronounced for CDC1551 at high nitrate levels.

## Discussion

The concept of morphological remodeling in response to diverse environmental signals has been extensively studied and observed in a variety of bacterial species, such as *Helicobacter pylori*^53^, *Campylobacter jejuni*^54^, or *Vibrio parahaemolyticus*^55^. The transition from helicoidal or rod-shaped to coccoid morphologies in these species has been described as an adaptive strategy that allows bacterial survival under changing environmental conditions. In particular, remodeling of their cell architecture represents a critical survival mechanism, allowing bacteria to effectively reduce antimicrobial influx and increase tolerance to various stresses^56^. This highlights the importance of physical adaptation to environmental pressures for prokaryotes. Morphological changes are frequently linked to metabolic remodeling accompanied by genome-wide transcriptional reprogramming^57,58^. Beyond global morphological adaptations, bacteria can also undergo intracellular architectural reorganization in response to external stimuli, notably through the formation of complex and extensive cytosolic compartments^7,59^. However, observations of this phenomenon have been studied mainly in environmental bacterial species, due to the complexity and diversity of external signals encountered in these niches. In contrast, intracellular pathogens also face a broad spectrum of stressors within the host and although similar remodeling of their internal architecture might be expected, this process remains comparatively underexplored. Importantly, processes leading to internal reorganization and compartmentalization in bacteria could represent new potential targets for antimicrobial strategies, which have not been visited so far.

In our study, we used cryo-electron tomography to investigate various strains of *Mycobacterium tuberculosis*, a clinically important human pathogen, and related mycobacteria at nanometer-scale resolution, uncovering many aspects of their internal cellular architecture. One of our most intriguing findings was the varying presence of intracytoplasmic membranes when comparing members of the MTBC, such as Mtb and *M. bovis* BCG with extensive MIM structures, with the non-pathogenic mycobacterial model organism *M. smegmatis* that lacked MIMs (see Figure 1, Supplementary Figure 2). This suggests that MIM formation may be a specific adaptation behavior of intracellular mycobacteria. We also show heterogeneity in MIM formation within MTBC members and individual Mtb strains, which raises questions about possible differences in the microevolution of selected Mtb and MTBC strains in terms of host-adaptative strategies. Cryo-ET on FIB-milled lamellae of intra-and extracellular Mtb revealed a variety of size, number and localization of MIMs within bacteria, indicating that these characteristics are likely not tightly regulated. We also re-examined published TEM images of Mtb, and we identified visible intracytoplasmic membranous structures, although they were not mentioned by the authors of the study^60^. Similarly, very recent work includes cryo-tomogram slices in which other intracytoplasmic membranous features are apparent^61^. These were also not further highlighted nor discussed by the authors, and we therefore cannot conclude whether the structures of the mentioned studies corresponded to the MIMs that we show and discuss in our study. Nonetheless, these observations highlight a potential broader distribution of internal compartmentalization features in pathogenic bacteria that has yet to be explored. Recently, cytoplasmic, filamentous, clustered tailocins have been characterized by cryo-ET in the pathogen *Listeria monocytogenes*^62^. Tailcins slightly resemble MIMs, however they largely differ in their organizational pattern within Listeria, their function related with bacterial growth, and their lack of a surrounding membrane^62^.

Our cryo-ET datasets allowed a detailed characterization of intracytoplasmic membranes including MIMs in *Mycobacterium tuberculosis*. While our findings suggest that MIMs originate from invaginations of the inner membrane, a process also observed in other bacterial phyla^63^, the mechanisms underlying ICM biogenesis and the incorporation of membrane proteins remain poorly understood. This has been a fundamental question in other ICM-forming bacteria, particularly in relation to thylakoids formation in cyanobacteria^7,64,65^. Their modulation by varying light intensities has been explored in detail, and their protein content has been resolved^7,66^, but the molecular basis of how such extensive membrane networks are dynamically modulated, stabilized and maintained with enzymatic activity remains elusive. It is important to note that while cryo-ET is increasingly used to gain insights into subcellular organization in microorganisms, it is a static technique that does not capture dynamic processes. Thus, it remains unclear in our settings whether the Mtb MIMs connected to the IM are in the process of being invaginated into the cytosol, or whether they instead fuse with the IM after being recruited from the cytosol. We hypothesize that the basal turnover of MIMs observed *in vitro* is exacerbated during intracellular infection, where multiple host-induced stresses could impact the pathogen and MIM dynamics (see Figure 2). We point out that an understanding of the biogenesis, dynamics and link with metabolism of ICM compartments in bacteria and particularly pathogens like Mtb could provide new potential targets for antimicrobial intervention.

In our study, we reveal that Mtb MIMs are associated with an enrichment of NarS, the histidine kinase of a two-component system involved in nitrate sensing^36^. During infection, Mtb often encounters hypoxic or near-anaerobic environments where the bacilli can switch to nitrate respiration, using nitrate as a terminal electron acceptor when oxygen is limited^33^. Expanding membrane surface area through intracytoplasmic membrane formation, particularly to accommodate nitrate-responsive proteins, may support this metabolic shift and support detoxification of reactive nitrogen species in Mtb during host infection. Similar processes have been described in ammonia-oxidizing bacteria like Nitrobacter sp., which form ICMs that host membrane-bound ammonia monooxygenase to perform enzymatic reactions allowing the oxidation of ammonia to nitrite^67^. In Mtb, the expression of the NarGHIJ and NarX nitrate reductases, as well as nitrate/nitrite transporters (NarK2, NarK3, NarU), is not directly induced by nitrate in the environment, but rather by hypoxia^32^. In our experimental conditions, where oxygen levels were kept at atmospheric levels, expression of these reductases and transporters remained unchanged when nitrate was supplemented. In contrast, *NarS* and its response regulator *NarL* were both upregulated, alongside strong induction of MIM formation (see Figure 4). This uncoupling suggests that NarL-mediated gene regulation may operate independently of classical nitrate reduction pathways and could involve co-regulation with other transcriptional regulators. Notably, previous studies have demonstrated that DosR, the central regulator of the response to hypoxic environments and nitric oxide, shares a partially overlapping regulon with NarL^36^. Supporting this uncoupling of NarSL function and nitrate reduction activity, no NarL-binding sites have been identified in the promoter regions of *narL*, *narGHIJ*, or *narK* genes^68^ in contrast with the clear regulatory roles of NarL homologs in *E. coli* and *Pseudomonas aeruginosa*^49,69–71^. In Mtb, the DosRST two-component system is well characterized and is known to be activated by environmental cues such as hypoxia and NO, both conditions present during host infection that lead together to triggering of Mtb dormancy^29,72–74^. In our settings, we observed increased expression of DosR and DosS upon addition of nitrate to the medium (see Supplementary Figure 9). However, whether this activation results from direct nitrate sensing or from NO generated via basal nitrate reduction remains unclear.

The Mtb strains used during this study emerged from unrelated epidemics in space and time^75,76^, and have been shown to differ in their immunogenicity in animal models^45,51,77^. We found that the same strains also exhibit strong differences in MIM content and exhibited largely different capacities inducing them. Furthermore, our study revealed that infection of host cells with MIM-induced Mtb led to lower levels of inflammation markers in the supernatant of both macrophages and primary dendritic cells compared to Mtb grown under control conditions. These observations suggest that formation of MIMs not only remodulates Mtb metabolism in response to nitrate in the environment but also influences host pathogen interactions during infection. This dual role of MIMs in Mtb places these structures as a main hub involved in environmental adaptation within the host. Furthermore, the immunogenic heterogeneity observed across Mtb strains and their sensitivity to MIM-inducing signals could broadly influence host responses and could be critical for optimizing treatment strategies. We note that whether the observed lower inflammation results from reduced expression of immunogenic bacterial proteins or from reduced host cell death and damage remains to be determined.

In conclusion, our study provides the first description of mycobacterial intracytoplasmic membranes with unprecedented ultrastructural insights into the inner organization of Mtb and other pathogenic mycobacteria and suggests that MIM formation is tightly linked to nitrate sensing and metabolic adaptation. These findings thereby also raise new questions about compartmentalization of metabolic processes in intracellular bacterial pathogens, which might be broadly distributed in the bacterial world. Our results also reveal new insights on the regulatory networks connecting nitrate sensing, nitrogen use and immune evasion in Mtb infection processes, and open yet unexplored avenues for assessing the full biological functions of MIMs, including their potential as putative anti-mycobacterial drug targets.

## Materials and methods

### Cell culture and cell lines

RAW264.7 macrophages (ATCC TIB-71) were cultured in Dulbecco’s modified Eagle’s medium (DMEM, Gibco 12077549) supplemented with 10% (v/v) heat-inactivated fetal bovine serum (Sigma-Aldrich) at 37 °C, 5% CO_2_ (referred to as “complete DMEM medium” for the rest of the manuscript) and were continuously monitored for mycoplasma contamination.

### Mycobacterial strains and classic culture conditions

All strains used in this study were either taken from our in house collection (Institut Pasteur Paris, France), or they were kindly provided by M. Guttierrez (The Francis Crick Institute, London UK). Mtb H37Rv, Mtb H37Rv ΔESX1, Mtb H37Rv ΔPDIM, Mtb CDC1551, Mtb Erdman, *Mycobacterium bovis* BCG, *Mycobacterium smegmatis* MC^2^1551 were cultured in Middlebrook 7H9 broth base (Sigma-Aldrich M0178) supplemented with 0.2% glycerol (Fischer Scientific), 10% albumin-dextrose-catalase (ADC; BD 211887), and 0.05% Tween-80 (Sigma-Aldrich), or on Middlebrook 7H11 agar base (Sigma-Aldrich M0428) supplemented with 10% Oleic Acid-ADC (OADC; BD 211886). When required, appropriate selection antibiotics were used: kanamycin at 25 µg.mL (Sigma-Aldrich K1876), hygromycinB at 50 µg.mL (Thermo Fischer 10453982), and NaNO_3_ (Sigma-Aldrich) was added to the cultures as described in the results section at 10 mM, 50 mM or 100 mM. Cultures were incubated at 37°C without CO_2_ and no agitation. Adaptions of culture conditions for cryo-electron microscopy experiments are described in a dedicated section.

### Macrophage infection with Mtb

RAW264.7 macrophages were seeded 24 hours prior to infection. Mycobacteria cultures in mid-log phase (0.8±0.2) were pelleted at 3500 g for 5 minutes and washed twice in PBS (Gibco). Pellets were resuspended in 10 mL of cell culture media and adjusted to the same OD_600nm_ and appropriate dilutions were made for infection. The bacterial inocula were added to cells and incubated for 2 hours of uptake at 37°C and 5% CO_2_. After 2 hours of uptake, infected cells were washed twice with PBS and incubated in complete medium supplemented with gentamicin (50 µg.mL) for 1 hour at 37°C 5% CO_2_ to kill remaining extracellular bacteria. Cells were washed twice in PBS and re-incubated with complete DMEM medium until appropriate timepoints. When required, infected cells were fixed for 1 hour at room temperature with 4% paraformaldehyde in PBS (Thermo Fischer). For all experiments in this study, we assumed an OD_600nm_ of 1.0 corresponds to 1.0x10^8^ bacteria per mL. For colony-forming units (CFU) assays, infected cells were lysed using 0.05% Triton-X100, and serial dilutions were plated on 7H11 agar plates, and incubated for 3 weeks at 37°C before counting colonies.

### Sample preparation for cryo-ET

Cryo-EM silicon grids (Quantifoil SiO_2_ film R2/2 Au 200 mesh) were placed on a 35 mm dish (Ibidi), plasma-cleaned for 45 seconds at 15mA and 0.38 mbar using PELCO easiGlow, and UV-sterilized before cell seeding.

For the investigation of phagocytosed pathogens, RAW264.7 macrophages were seeded on top of the freshly glow-discharged EM grids at a density of 2x10^5^ cells for one 35 mm dish, giving a final concentration of 1-3 cells per grid-square. The montage (dish – EM grids – cells) was incubated 24 hours at 37°C and 5% CO_2_ in complete DMEM until cell adherence to the grids, before infection with Mtb. The mycobacteria were cultured in 7H9 – 10% ADC – 0.2% glycerol *without* Tween-80. Cultures in mid-logarithmic phase (OD_600nm_ 0.8±0.2) were pelleted for 5 minutes at 3500 g and washed twice with PBS. Pellets were vortexed with glass beads to break large clumps before resuspension in 10 mL of complete DMEM, and the remaining aggregates were allowed to sediment for 5 minutes at room temperature in a still tube. The top 7mL of the “supernatant” was filtered through a 10 µm strainer (pluriStrainer, Dutscher) to remove smaller aggregates and the OD_600nm_ of the filtrate was measured. Cultures were adjusted to the same OD_600nm_ and appropriate dilutions were made to infect cells at a final MOI of 1 or 5. After 2 hours of uptake, cells were washed 3 times with PBS to remove extracellular bacteria, resuspended in complete DMEM, and incubated at 37°C and 5% CO_2_ for 4 hours or 24 hours. At appropriate times post-infection, infected cells were washed twice with PBS and inactivated for 30 minutes at room temperature in 2% PFA (Electron Microscopy Science), 0.05% glutaraldehyde (Sigma-Aldrich), 0.2M HEPES (Gibco), then 30 minutes in 4% PFA, 0.1% glutaraldehyde and 0.2M HEPES. After fixation, grids were resuspended in PBS and immediately vitrified.

For cryo-ET experiments on bacterial suspension only, the mycobacteria were prepared and inactivated as described above omitting the step adding them to the macrophages.

### Plunge freezing/vitrification of infected cells and bacterial suspension

The EM grids with bacteria alone or macrophages containing bacteria were transferred to a Leica Microsystems EM GP2 automated plunge freezer and adjusted to the chamber conditions (18°C 98% humidity) for 15 seconds. The grids were then back-side blotted with a filter paper for 8 seconds. For vitrification of mycobacteria, 3 μL of inactivated bacterial suspension were pipetted on Quantifoil SiO_2_ R2/2 grids freshly glow discharged for 90 seconds at 15 mA and 0.38 mbar (PELCO easiGlow). Grids were transferred to the EM GP2 plunge freezer, adjusted to the chamber conditions (18°C 98% humidity) for 15 seconds and excess liquid was back blotted for 4 seconds. Grids were plunge frozen in liquid ethane at liquid nitrogen temperature (-180°C) and stored in sealed boxes in liquid nitrogen until further use.

### Cryo-FIB milling

Plunge frozen grids were clipped with c-clips (Thermo Fisher Scientific 1036171) into cryo-FIB autogrids (Thermo Fisher Scientific 1205101) and mounted into a 35° pre-tilted FIB-shuttle (Thermo Fisher Scientific 1213330). Samples were transferred to the cryo stage of a dual beam Aquilos2 cryo FIB-SEM instrument (Thermo Fischer Scientific), using its cryo-transfer system. SEM maps of the grids were acquired using the Maps v3.29 (Thermo Fischer Scientific) software to place targeted milling positions.

After selecting milling positions, grids were coated with an organo-metallic platinum layer using a gas injection system (GIS) for 1 min before milling. Lamellae were automatically milled to 200 nm with a stepwise reducing current (1 nA–10 pA) using automated cryo-FIB milling software AutoTEM (v.2.4.3 Thermo Fisher Scientific) with a 10° milling angle. First, stress relief cuts (0.6 μm × 8 μm × 7 μm) were made 4 μm away from the lamellae, which were then set to ∼12 μm width, using a milling current of 1 nA. A step of rough milling was then performed to reduce the lamella to 1 μm thickness using a rectangular pattern on both sides of the lamellae simultaneously with a milling current of 1 nA. Medium milling further reduced the lamella to 500 nm using the cleaning cross-section pattern on each side alternately with a milling current of 0.1 nA. Fine milling thinned the lamella to 300 nm using the same pattern as medium milling with a milling current of 50 pA. Finally, an automated thinning step was done with a current of 10pA to reduce the thickness of the lamella to ∼200nm. Lamellae were then manually polished to ∼100 nm using a 10pA current. SEM imaging was performed at 2-5kV and 13pA simultaneously to FIB-milling, to monitor the state of the lamellae. Grids were unloaded and stored in liquid nitrogen until data collection.

### Cryo-electron tomography data collection

Cryo-ET datasets were collected on a cold-Field Emission Gun Titan Krios (Thermo Fischer Scientific) transmission electron microscope operated at an acceleration voltage of 300kV, equipped with a Falcon 4i direct electron detection camera (Thermo Fischer Scientific) and a SelectrisX energy filter (Thermo Fischer Scientific), operated in counting mode. Tilt series (TS) acquisitions were done using the Tomography5 software v5.6 (Thermo Fischer Scientific). The microscope was set to nanoprobe mode for data collection, the energy filter slit width to 10eV, the C2 lens to 70 μm and the objective aperture to 100 μm. TS were acquired with a dose-symmetric scheme^78^ using 3° angular increments between +70° and -50°, starting with a 10° tilt to compensate for the lamellae pre-tilt. Nominal magnifications were varied between the acquisitions and ranged from 26 000 x, 42 000 x to 53 000 x, corresponding to a pixel size of 4.767 Å, 3.101 Å, and 2.453 Å respectively. The defocus value used ranged from -3 to -5 μm and the total dose applied was set to 140e^-^/Å^2^. Frames were saved in eer format, and motion correction was done on-the-fly by the TomoLive software^79^ (Thermo Fischer Scientific) with an algorithm based on UCSF MotionCor2^80^. The cold-FEG was flashed before every acquisition.

In total, 130 TS were acquired for Mtb H37Rv, 31 TS for Mtb H37Rv ΔESX1, 13 TS for Mtb H37Rv ΔPDIM, 131 TS for Mtb CDC1551, 12 TS for Mtb Erdman, 16 TS for *M. bovis* BCG Pasteur, and 40 TS for *M. smegmatis* MC^2^155. For samples of infected cells, a total of 19 TS of Mtb H37Rv-infected macrophages were acquired, 27 TS of Mtb CDC1551-infected macrophages, 3 TS of Mtb Erdman-infected macrophages and 9 TS of *M.bovis* BCG Pasteur-infected macrophages. Cryo-ET data of infected cells were acquired at 4 hours or 24 hours post-infection.

### Tilt series alignment and reconstruction

Motion-corrected tilt series were manually inspected to remove high-angle bad tilts when necessary and aligned through patch-tracking on IMOD. Aligned TS were reconstructed on IMOD^81^ using weighted-back projection and SIRT-like filter with 10 iterations. Tomograms were inspected for their alignment, thickness and overall quality. For selected tomograms, 2D contrast transfer function (CTF) was calculated with ctfplotter and the correction was applied with phase-flipping using ctfphaseflip^82^. Tomograms were reconstructed to a pixel size of ∼10 Å corresponding to a binning of 2 to 4 depending on the original pixel size of each dataset. Reconstructed tomograms were further filtered using isotropic reconstruction software IsoNet^83^.

### Tomogram segmentation and template matching

Selected (reconstructed and isonet-corrected) tomograms were used as input for automated segmentation using MemBrain-seg^84^ segmentation tool with the v10_alpha pre-trained model. Resulting segmentations were manually curated on Napari0.5.6 for further refinement and removal of false positive detections. For visualization purposes and movie-making, cleaned segmentations were imported into UCSF ChimeraX-1.9^85^.

Cross-correlation-based template matching of mycobacterial and host macrophage ribosomes was performed using the PyTom standalone python package^86^. Template matching was run with EM maps of the Mtb 70S initiation complex (EMDB-23974) and the mouse 80S ribosome (EMDB-30432), down sampled to the tomogram’s pixel size. A 350Å diameter mask was applied to the cross-correlation. The candidates cross-correlation cut-off was set to 0.25 and the maximum number of particles to extract was set to 10 000 within the cut-off limit. The obtained matched candidates were added on their corresponding tomograms using the ArtiaX toolbox^87^ of UCSF ChimeraX, and individual particles were manually inspected to ensure a correct correlation between a particle and its corresponding density on the tomogram. False positives were removed by sliding the CCLmax value upwards and remaining false positives were manually removed.

### Cryo-ET data analysis

MIMs were manually counted in all tomograms obtained. Measurements of MIM length and mycobacteria membrane thickness were done using IMOD and Fiji^88^.

### Sample preparation for Mass Spectrometry proteomics analysis

Cultures of Mtb H37Rv, Mtb CDC1551 and *M. smegmatis* MC^2^155 were grown in Middlebrook 7H9 Broth Base supplemented with 10% ADC, 0.2% glycerol and *without* Tween-80. Five biological replicates of each strain were prepared. To enrich MIMs for MS analysis, the following protocol was used with some adaptations to our BSL3 settings^89^. Bacterial cultures at OD_600nm_ 0.8±0.2 were centrifuged at 3500 g for 5 minutes at 4°C and washed in PBS twice. Pellets were resuspended in 10 mL of PBS supplemented with a tablet of cOmplete EDTA-free protease inhibitor cocktail (Sigma) and 0.1mm zirconium beads (BioSpec 11079101z) before transferring into small Nalgene containers.

Containers were mounted on a TissueLyser2 (QIAGEN), and suspensions were mechanically lysed in at maximum speed (30Hz), with three rounds of 1 minute lysis – 1 minute incubation on ice. Lysates were retrieved and centrifuged at low speed (200 g for 7 minutes) to remove beads and bacteria debris. Supernatants were double filtrated through 0.22 μm filters and centrifuged 40 minutes at 4°C 10 000 g to separate the mycomembrane from the inner membrane and cytoplasmic components. Supernatants were recovered and centrifuged again 30 minutes at 4°C, 27 000 g. Supernatants were recovered and ultracentrifuged for 40 minutes at 4°C, 100 000 g. Pellets were recovered and resuspended in PBS. Different density gradient solutions (5% – 20% – 30% – 40% – 50% – 60%) were prepared by diluting a 60% Opti-prep^TM^ stock solution (60% Iodixanol; Sigma-Aldrich) with PBS. Pellets resuspended in PBS were added at the top of the tubes above the top layer of the density gradient. The gradient was ultracentrifuged for 2 h at 100 000 *g* at 4 °C (Beckman SW 60 Ti rotor). After ultracentrifugation, fractions of 0.4 mL were collected from the top of the tube. The protein concentration was measured with the microBCA kit (Pierce).

For each biological replicate, 20 micrograms of protein from each bacterial strain were used. Proteins were precipitated using TCA/acetone and subsequently resuspended in a solution containing 5% SDS and 50 mM TEAB (T7408 Sigma-Aldrich). Digestion was performed using S-Trap™ micro spin columns (C02-micro-40, Protifi) following the manufacturer’s instructions, with minor modifications: iodoacetamide (IAA) replaced MMTS as the alkylating agent, and 2 micrograms of trypsin (V5111, Promega) were used per sample in 20 µL of 50 mM TEAB (pH 8.5), incubated overnight at 37°C. Eluted peptides were pooled, dried using a SpeedVac, and resuspended in 0.1% formic acid (FA) and 2% acetonitrile (ACN) in HPLC-grade water. Samples were then desalted using C18 stage tips prior to LC-MS/MS analysis, which was conducted using a timsTOF Ultra mass spectrometer equipped with a CaptiveSpray source (Bruker) and coupled with a nanoElute® 2 nanoflow liquid chromatography system (Bruker). Samples were loaded at 800.0 bar on a PepSep® ULTRA (25 cm x 75 µm x 1.5 µm) C18 HPLC column (Bruker) and equilibrated in 98% solvent A (H2O, 0.1% FA) and 2% solvent B (ACN, 0.1% FA). Peptides were eluted using a 2 to 4% gradient of solvent B during 1 min, then a 4 to 20% gradient of solvent B during 19 min, followed by a 20 to 30% gradient of solvent B during 5 min and finally a 30 to 95% gradient of solvent B during 1 min all at 250 nl/minute flow rate.

The instrument method for the timsTOF Ultra was set up in a data-independent acquisition (DIA) scan mode that uses parallel accumulation-serial fragmentation (PASEF®) technology (dia-PASEF®). The mass range for MS scans was set up to 100-1700 m/z, whereas the mobility (1/K0) range was set up to 0.64-1.45 Vs/cm² with a 100.0 ms ramp time. The number of MS/MS windows was set up to 24 with a width of 25 Da spanning from 400.0 to 1000.0 Da.

Raw data were analyzed using Spectronaut® v19 (Biognosys). All searches were performed with oxidation of methionine and protein N-terminal acetylation as variable modifications and cysteine carbamidomethylation as fixed modification. Trypsin was selected as protease allowing for up to two missed cleavages. The maximum and minimum peptide length were set to 52 and 7 amino acids, respectively. The false discovery rate (FDR) for peptide and protein identification was set to 0.01.

For the identification of differentially abundant proteins between conditions, a statistical analysis pipeline was implemented from the intensities quantified by the Spectronaut software using an R script. First, proteins were filtered to retain only high-confidence identifications. Reverse hits (potentially misidentified proteins), potential contaminants, and proteins “Only identified by site” were removed from the dataset. Only proteins with at least one unique peptide (peptides not common to other proteins in the FASTA file) were considered for quantification. Proteins were retained only if they had at least 2 quantified intensity values in a condition. Raw intensity values were log-transformed (log2) and normalized within each condition. The normalization procedure centered the values on the mean of the medians of intensities in each sample of the condition^90^. Proteins present in one condition but absent in another were set aside. For the remaining proteins with missing values, imputation was performed using the impute.mle function from the R package imp4p^91^. Differential abundance analysis was conducted using a limma t-test^92^. The resulting p-values were adjusted using an adaptive Benjamini-Hochberg correction through the adjust.p function of the cp4p R package^93^. The proportion of true null hypotheses was estimated using the pounds method^94^. Proteins were considered significantly differentially abundant if they met 2 criteria: an adjusted p-value below 0.01 (corresponding to a false discovery rate of 1%) and an absolute log2(fold-change) greater than 2.

Presence of transmembrane domains was predicted using deepTMHMM^95^. GO analysis was performed using DAVID^96,97^. The UniProt accession numbers of each protein were used as input. Volcano plots and dot plots were performed on R studio v4.4.1.

### Quantitative Real-Time Polymerase Chain Reaction (qRT-PCR)

Mtb H37Rv and Mtb CDC1551 were cultured as described above until midlogarithmic phase. Cultures were diluted to OD_600nm_ 0.05 in 50mL with or without 100mM NaNO_3_ and grown until OD_600nm_ reached 0.8±0.2 before proceeding to mycobacterial RNA extraction. Cultures were centrifuged 10 minutes at 3500 g, 4°C and pellets were resuspended in 1mL of TRIzol® reagent (Thermo Fischer 15596018). Suspensions were transferred into 2mL Lysing Matrix B tubes containing 0.1 mm silica beads (MP Biomedicals) and mechanically lysed using a Fast-Prep machine. Tubes were recovered and centrifuged at 10 000 g, 4°C, for 10 minutes to remove beads and bacteria debris.

200 μL of chloroform was added to the supernatant of each tube and shaken vigorously for 10 seconds before centrifuging the tubes at 10 000 g, 4°C, 15 minutes. The aqueous phase was recovered in new RNAse-free tubes and an equivalent volume of isopropanol was added before mixing the tubes and storing overnight at -20°C. Tubes were then centrifuged at 13 000 rpm, 4°C, for 20 minutes and supernatant was carefully removed. The pellet was washed with 500 μL of cold 70% ethanol and centrifuged 15 minutes, 13 000 rpm, 4°C. Ethanol was removed and pellets were dried before resuspension in 100 μL of RNAse-free water. The extracted RNA was treated with DNAse (TURBO DNAse-free kit, Ambion Life Technologies) according to manufacturer’s recommendation. RNA samples were stored at -80°C until further use. RNA was reverse transcripted with the SuperScript III reverse transcription kit (Thermo Fisher) and cDNA was stored at -20°C until use. cDNA samples were used for quantitative real-time amplification of target genes, using the SYBR green qPCR mastermix (Thermo Fischer). In each sample, the amount of amplified cDNA of each target gene was normalized to the quantity of the mycobacterial housekeeping gene rpoB, in the same strain and condition. Oligonucleotides sequences used for amplification of cDNA targets are listed in Supplementary Table 4.

The relative change in target gene expression between strains and conditions was calculated with the threshold cycle ΔΔCt method^98^. At least 3 independent experiments were performed on 2 biological replicates.

### Cell infection and Enzyme Linked ImmunoSorbent Assay (ELISA)

Bone Marrow-derived Dendritic Cells (BM-DCs) were generated from 8-week-old C57BL/6J (H-2^b^) female mice (Janvier) in RPMI 1640-GlutaMax medium (Gibco), complemented with 10% fetal bovine serum (Gibco), 1% of MEM non-essential amino acids (Invitrogen, Life Technologies), 5 x 10^-5^ M β-mercaptoethanol (Invitrogen, Life Technologies) and 25 ng/mL of recombinant mouse GM-CSF (Gibco). Differentiated cells were used at day 7 of culture for infection with mycobacteria. BM-DCs were collected, washed and seeded in 48-well plate (TPP) at 2.5 x 10^5^ cells/well in RPMI 1640-GlutaMax medium containing 10% FBS. Four hours later, cells were infected with *Mtb* H37Rv or *Mtb* CDC1551 strain grown with or without 100mM NaNO_3_ as described in the corresponding section at various M.O.I.. After 24 hours and 48 hours of infection at 37°C and 5% CO_2_, supernatants were collected, filtered and key pro-and anti-inflammatory cytokines were quantified by ELISA using R&D Biosystems DuoSet kits according to manufacturer’s recommendations. Experiments were performed in biological duplicates and technical duplicates. RAW264.7 macrophages were seeded 24 hours before infection in 48-well plates at 2.5x10^5^ cells/well in complete DMEM medium. Cells were infected as described above.

### Figures and statistics

All figures were done on R Studio v4.4.1 (Posit team (2024). RStudio: Integrated Development Environment for R. Posit Software, PBC, Boston, MA. URL http://www.posit.co/.) and GraphPad Prism 10.2.2 (GraphPad Software for Mac, Boston, Massachusetts USA, www.graphpad.com). Statistics were performed on GraphPad Prism, with a *p*-value considered significant if *p* < 0.05 with * < 0.05, ** < 0.01, *** < 0.001 and **** < 0.0001. Figures were put together on Adobe Illustrator v29.1, and some elements of figures (Fig. 1A, 3A) have been created on BioRender (biorender.com).

## Acknowledgments

We thank Sandrine Schmutz (Flow Cytometry Platform) of C2RT for technical assistance during cell sorting, Mohamad Harastani (Image Analysis Hub), Eduard Baquero-Salazar and Anna Sartori-Rupp (Nanoimaging Core Facility), Anastasia D Gazi (Ultrastructural BioImaging facility, UBI) for help on cryo-tomograms analysis and helpful discussions, and the UBI facility for trainings provided on plunge-freezing and negative staining TEM. We also thank Maximiliano Gutierrez (The Francis Crick Institute, London, UK) for sharing tools. We thank the current members of the DIHP unit especially Camila Valenzuela, Elif Begum Gokerkucuk and Arthur Lensen for helpful discussions, as well as the current members of the PMI unit, especially Wafa Frigui, Cécile Tillier and Mickael Orgeur. C.K wishes to thank Priscille Brodin (Institut Pasteur, Lille, France), Léa Swistak (EMBL Heidelberg, Germany), Gautham Sankara Nayarana (Institut Pasteur, Paris, France), and Cyril Anjou (Centre de Recherche Saint-Antoine, Paris, France) for advice, feedback, and support.

## Author contributions

C.K and J.E conceived the study with input from R.B. F.S provided the bone-marrow derived dendritic cells isolated from mice. D.M prepared and ran samples for mass spectrometry under the supervision of M.M, and Q.G.G performed the analysis. S.T and M.V provided advice for cryo-ET acquisitions. C.K designed, performed, and analyzed all other experiments. C.K and J.E wrote the initial draft of the manuscript. J.E and R.B secured funding. All authors participated in proofreading and correcting the manuscript.

## Data availability

The mass spectrometry raw data has been submitted to PRIDE with the following reference number: 1-20250915-131742-116390100.

