## Supplementary for "Nitrate-responsive Mycobacterial Intracytoplasmic Membranes dampen Inflammation during *Mycobacterium tuberculosis* Infection"

#### Supplementary materials and methods

##### Generation of EGFP-Galectin3 RAW264.7 macrophage stable cell line

A RAW264.7 EGFP-Galectin3 stable cell line was generated using the sleeping beauty transposon system [1]. Briefly, the pSBbi EGFP-Galectin 3 transposon-containing plasmid was co-transfected with the vector encoding the SB100X transposase (pCMV(CAT)T7-SB100X (Addgene #34879) into host cells, using the JetPrime transfection reagent (PolyPlus 101000015). After 48 hours, cells were washed, and medium was exchanged with fresh complete DMEM medium supplemented with 3µg.mL Puromycin for selection. Transfected cells were selected for 7 days before expansion. Pools of EGFP-positive cells were sorted into sub-categories depending on EGFP expression levels using a BD FACSAria III Cell Sorter (BD Biosciences). A polyclonal pool of sorted cells was selected for further experiments.

##### Sample preparation for Cryo-ET of *Mycobacterium smegmatis*

Bacterial suspension of *M. smegmatis* MC$^2$155 was prepared as described in the main methods section for other mycobacterial species, omitting the step of inactivation.

##### Cryo-Correlative Light Electron Microscopy

For precise targeting of intracellular Mtb in various niches, EM grids with infected EGFP-Galectin3 RAW264.7 macrophages were imaged in fluorescence light microscopy using a Leica THUNDER Imager EM Cryo-CLEM (Leica Microsystems). A global map of the grid was acquired for later correlation and precise targeting for FIB-milling. Alternatively, precise regions of interest were also located with the iFLM Correlative System v1.4.0 (Thermo Fischer Scientific) of the Aquilos2 instrument.

##### Negative staining TEM microscopy

Samples obtained from density gradient ultracentrifugation as described in the corresponding main Methods section were used for negative staining TEM microscopy.

4ul of MiM-enriched fraction were adsorbed to freshly glow-discharged carbon-coated copper grids (CF200-CU-50, Electron Microscopy Science) for 30 seconds, washed three times with milli-Q water and stained for 30 seconds in 2% Uranyl Acetate. Grids were imaged at room temperature in a FEI Tecnai T12 transmission electron microscope operated at 120kV.

**pH measurements of mycobacterial cultures**

*Mycobacterium tuberculosis* strains H37Rv and CDC1551 were cultured as described in the main Methods section. Sodium Nitrate(100mM) was added to the cultures at day 0. At designated timepoints, 0.5 mL aliquots were collected to Eppendorf tubes and pH was measured using mQuant pH indicator strips (Sigma). pH was determined while the strip was still humid, as per the manufacturer's recommendation.

**Ethidium Bromide accumulation assay**

Mtb H37Rv was cultured with or without Sodium Nitrate (100mM) as described in the main Methods section. Cultures were harvested in exponential phase ($OD_{600nm}$ 0.8±0.2) and washed twice with PBS. Pellets were resuspended in PBS supplemented with 0.4% glucose (Gibco) and incubated for 20 minutes at room temperature. Aliquots of 100ul of bacterial suspension were added to the wells of 96-well plates, and ethidium bromide (Eurobio Scientific) was added at concentrations that ranged between 0.01$\mu$g/mL to 8$\mu$g.ml. Verapamil (Sigma) was added to appropriate wells at 70$\mu$g/mL. Fluorescence was measured using a FLUOStar plate reader using the 530 nm band-pass and the 585 nm high-pass filters as the excitation and detection wavelengths, respectively. Fluorescence data was acquired every minute for one hour at 37°C. Negative controls containing PBS-0.4% glucose were included. Experiment was performed independently in two biological replicates.

**AlphaFold binding predictions**

Binding predictions of Mtb proteins to the narS-narL intergenic region were performed using AlphaFold3[2]. Amino acid sequences of corresponding proteins and the nucleotide sequence of the narS-narL intergenic region were obtained from whole genome sequencing of our Mtb strains. Binding prediction probabilities are represented by the iPTM score, used to evaluate the confidence of each computed interaction.

#### Supplementary Figures legends

**Supplementary Fig. 1: Variety of internal features found in mycobacteria imaged with cryo-ET.** In most of our cryo-tomograms, we observe the dense ribosome-free nucleoid region surrounded by a higher density of ribosomes (A-E). We also observed intracellular vesicles of different shapes and sizes (A, C). Granules of α-glucans were consistently found in all strains imaged and appeared to localize predominantly toward the cell poles (B, D). Some of our cryo-tomograms show bacilli undergoing division, with the septum clearly visible (B, E), which in some cases appeared deformed (E). Additionally, in *Mycobacterium smegmatis* exclusively, we observed fiber-like densities oriented parallel to the cell body, resembling the architecture of bacterial cytoskeletal proteins (D). Images represented here are slices from tomograms acquired at magnification 42 000 x. Scale bar: 100nm.

**Supplementary Fig. 2: Montage of MIMs found in mycobacteria.** Crops of tomographic slices of Mtb H37Rv, CDC1551 and *M. bovis* BCG Pasteur. White arrowheads show individual MIMs. MIMs were found in close proximity with the IM as well as in the middle of the cytoplasm. They exhibited various degrees of membrane curvature. Scale bar: 50nm.

**Supplementary Fig. 3: MIMs orientation and localization within the cell. A.** MIMs are visible in our cryo-tomograms in the XZ plane. By rotating 90° the orientation, we can observe the side view of the MIM as illustrated in the XZ plane schematics and in Fig. 1C. Rotating another 90° in the other orientation would allow us to see the flat side of the MIM in the YZ plane. **B.** Preferred orientation of MIMs in mycobacteria, elongated along the X or Y axis. Some MIMs were too curved to determine a clear orientation. **C.** Sub-cellular localization of MIMs in mycobacteria.

**Supplementary Fig. 4: MIM formation and intracellular niche. A.** A stable cell line of EGFP-Galectin3 RAW264.7 macrophages was generated and infected with Mtb DsRed. Rupture of the phagosomal membrane induces a strong recruitment of EGFP-galectin3, allowing the specific targeting of phagosomal rupture. Images were obtained with a Leica widefield microscope equipped with a cryo-stage and a cryo-pump. **B.** Quantifications of

MIM-positive Mtb found in the phagosome or in the cytosol of infected macrophages. **C.** Graphic representation of MIM formation.

**Supplementary Fig. 5:** Negative stain transmission electron microscopy of MIM-containing fraction. Scale bar 50nm.

**Supplementary Fig. 6: Proteomics of MIM-containing fraction and putative involvement of DosRST in the regulation of NarS/NarL TCS. A.** Volcano plot of the 938 proteins identified after filters that selected only proteins present in ≥4 of 5 replicates and supported by >2 unique peptides in ≥n–1 replicates. Log2(Fold Change) thresholds are set to 1 and -1 ; p-value threshold is 0.05. **B**. Localization of Nar proteins identified in all 5 biological replicates by mass spectrometry and represented in the volcano plot. **C.** Genomic organization of the NarSL two-component system and the predicted domains of NarS and NarL. **D.** AlphaFold3 predicted interaction models between NarL and DevR and their co-binding to the NarSL intergenic region.

**Supplementary Table 1:** Proteins exclusively found in H37Rv (A) or CDC1551 (B) across all biological replicates.

**Supplementary Table 2: H37Rv MIM-associated proteome.** List of all proteins found in the H37Rv MIM-containing fraction, present in at least 4/5 replicates, with at least 2 unique peptides in at least n-1 of the replicates where said proteins were found, and with p-value < 0.05 and Log2(Fold-Change) > 1. Protein names are displayed with their description (fasta header), Log2(Fold-Change) and -log10(p-value).

**Supplementary Table 3: CDC1551-enriched proteins.** List of all proteins found in the MIM-less fraction of CDC1551, present in at least 4/5 replicates, with at least 2 unique peptides in at least n-1 of the replicates where said proteins were found, and with p-value < 0.05 and Log2(Fold-Change) < -1, corresponding to significantly CC1551-enriched proteins. Protein names are displayed with their description (fasta header), Log2(Fold-Change) and -log10(p-value).

**Supplementary Fig. 7: Tomographic slices of Mtb CDC1551 cultured in 100mM $NaNO_3$.** Black arrowheads show MIMs formed in Mtb CDC1551 after growth in nitrate. Slice number is displayed on top left corner. Scale bar: 100nm.

**Supplementary Fig. 8: pH and membrane permeability are not compromised by nitrate addition in the growth medium. A.** pH measurements in liquid culture containing Mtb H37Rv, CDC1551, or no bacteria, in the presence or absence of 100mM $NO_3$ overtime. Experiment was performed in two biological replicates. Representative image of pH measurements by pH strips (+/- 0.3) at day 7 in all cultures. **B.** Ethidium Bromide accumulation assay in Mtb H37Rv cultured with or without 100mM NO3 and associated area under the curve (AUC). Individual values, means and SD are shown. One-way ANOVA with Tukey's multiple comparisons test was performed. * indicates p-value < 0.05, ** <0.01, *** <0.001 and **** <0.0001. ns: non-significant.

**Supplementary Fig.9: Expression of DosRST in H37Rv and CDC1551 in the presence or absence of 100mM $NO_3$ in the growth medium**. Gene expression was compared by qRT-PCR between H37Rv or CDC1551 in the two conditions. Ct values were normalized to the expression of the housekeeping gene rpoB in their respective strain and in each condition. Experiment was performed on 2 biological replicates. Mean and SD are shown. One sample t-tests were used with comparison of the fold change to 1. * indicates p-value < 0.05, ** <0.01, *** <0.001 and **** <0.0001. ns: non-significant.

**Supplementary Fig. 10: Induction of MIMs in Mtb limits the inflammation levels in infected cells. A-B.** Cytokine quantifications by ELISA assay. Results are expressed as the relative percentage change in TNF$\alpha$ and IL-10 secretion by macrophages (A) or Bone Marrow-derived Dendritic Cells (B) infected with Mtb grown in the presence of 100 mM $NaNO_3$ ("$H37RV_{100}$", "$CDC1551_{100}$"), compared with Mtb grown under control conditions (without $NaNO_3$). Individual values, mean and SD are shown. Experiments were performed on biological duplicates. One sample t-tests were used with comparison to 0. * indicates p value < 0.05, ** <0.01, *** <0.001. ns: non-significant.

**Supplementary Movie 1: Cryo-electron tomogram of *Mycobacterium tuberculosis* H37Rv.** Segmented features show the mycomembrane, peptidoglycan and inner membrane. Scale bar: 100nm.

**Supplementary Movie 2: Cryo-electron tomogram of *Mycobacterium smegmatis* MC[2]155.** Segmented features show the mycomembrane, peptidoglycan, inner membrane and ribosomes. Scale bar: 100nm.

### Supplementary Figure 1

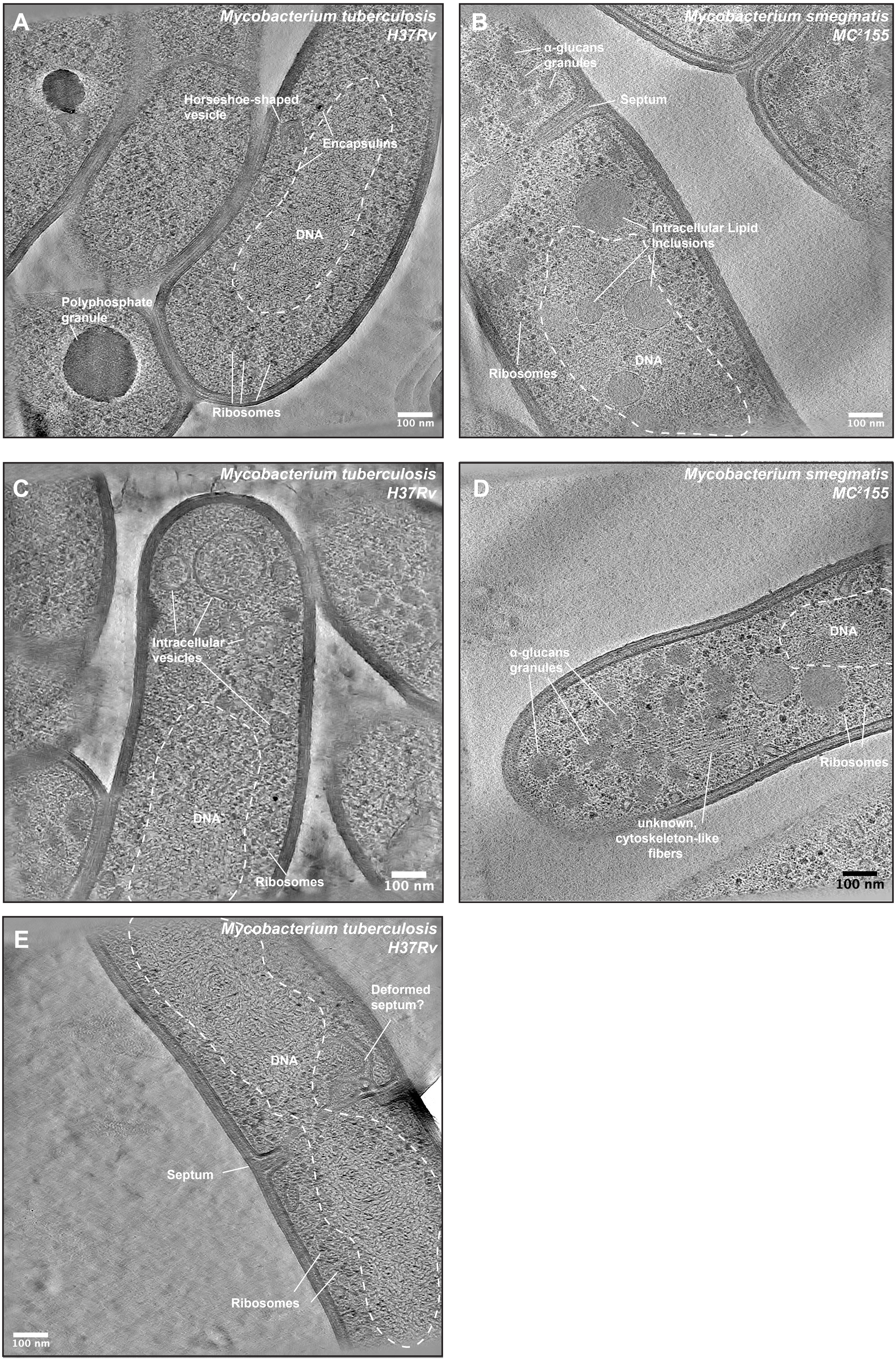

Supplementary Figure 2

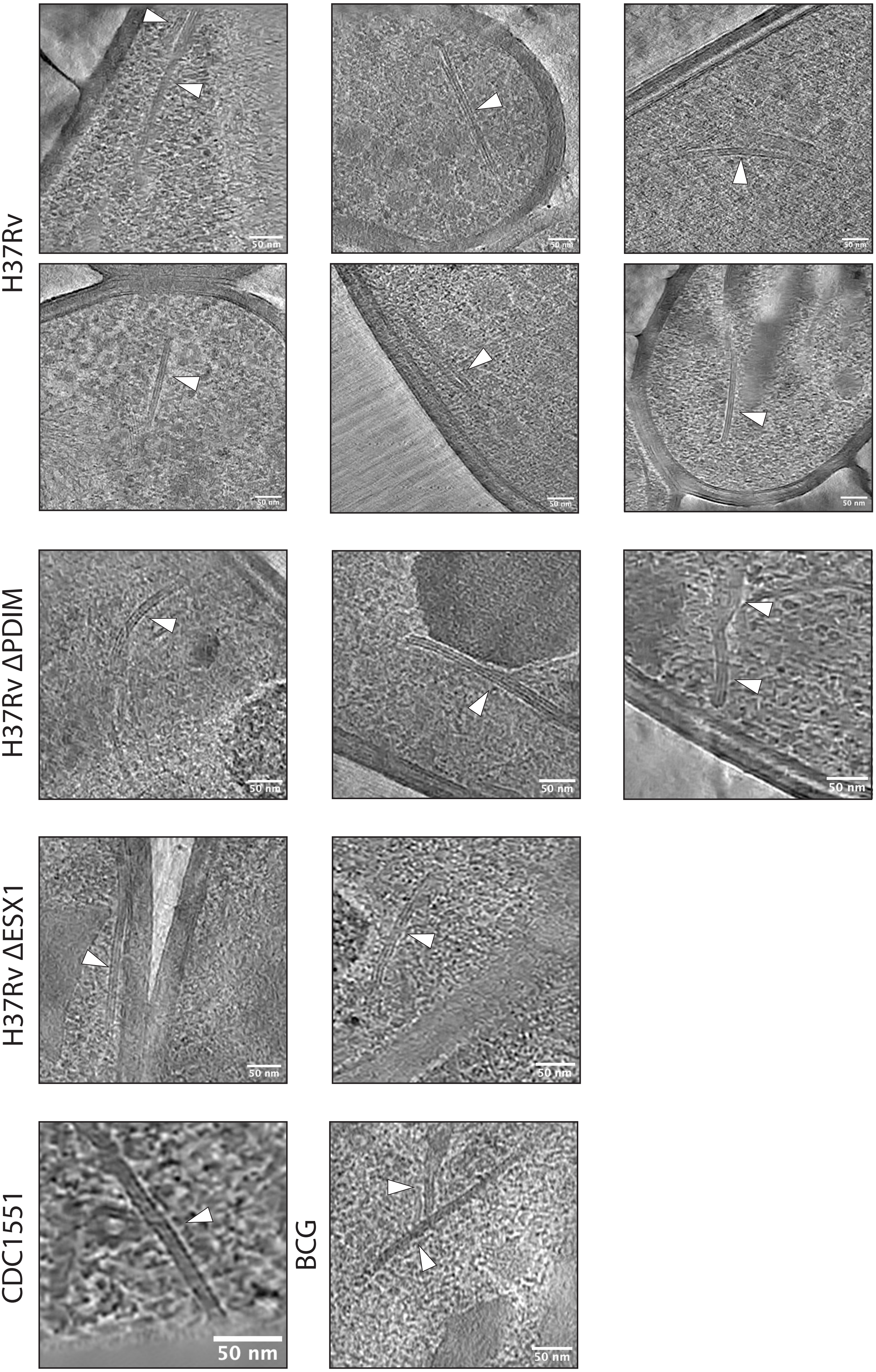

### Supplementary Figure 3

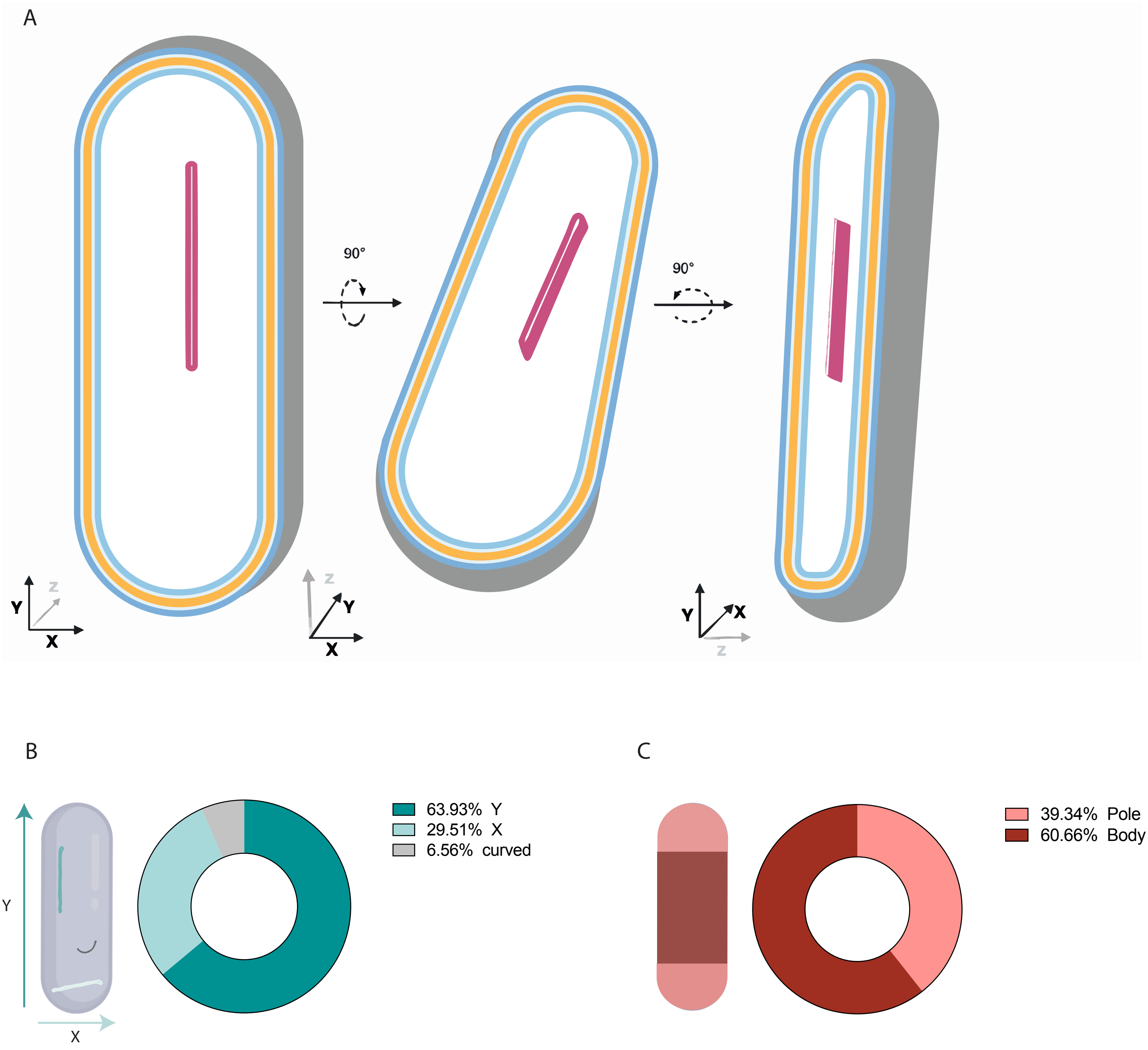

### Supplementary Figure 4

A

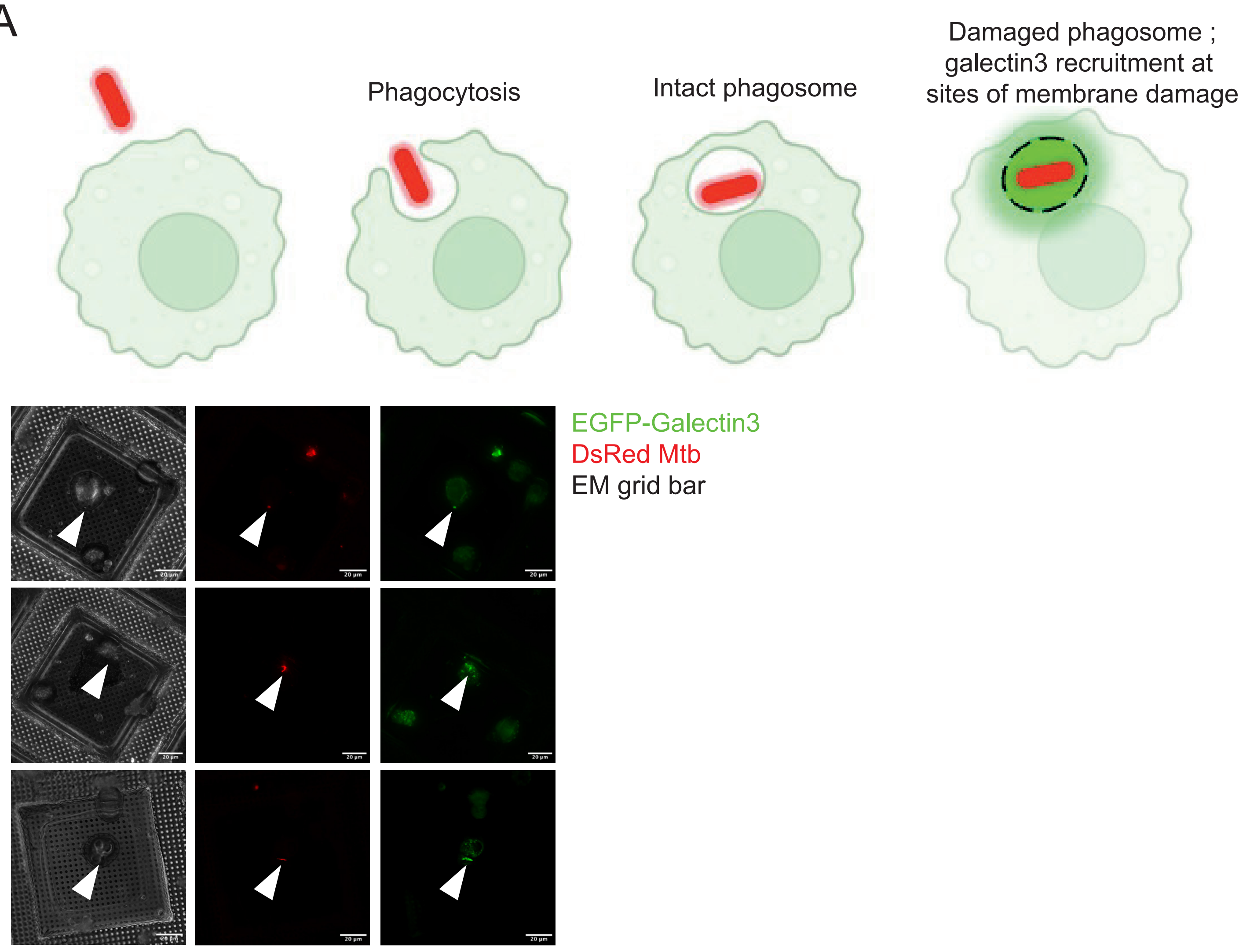

B

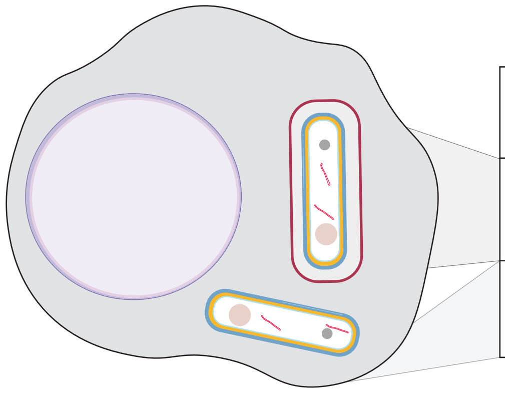

| | Mtb H37Rv | Mtb H37Rv |
|---|---|---|
| MIM-positive bacilli in phagosome | 0% | 60% |
| MIM-positive bacilli in cytosol | 100% | 40% |

C

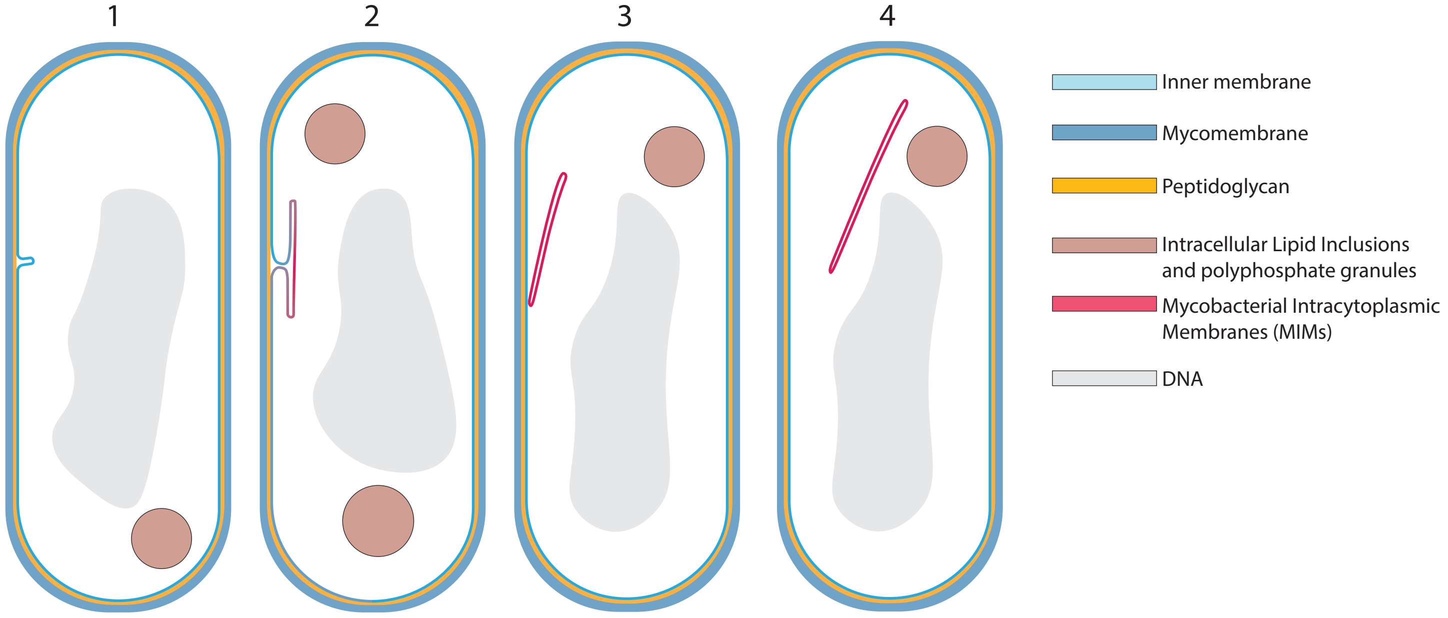

Supplementary Figure 5

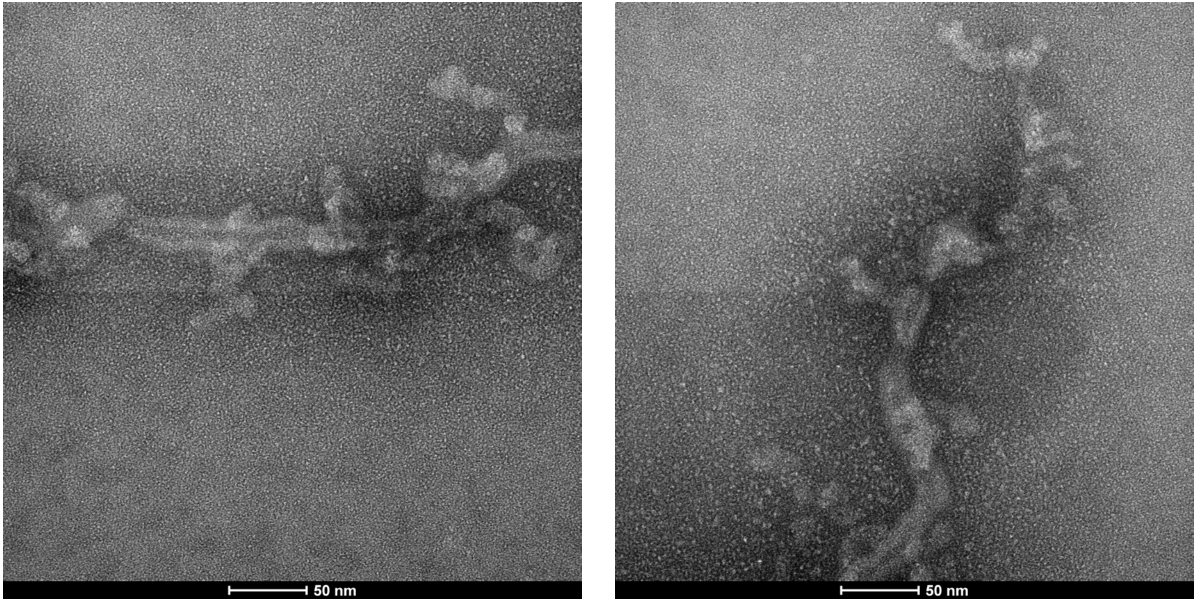

Supplementary Figure 6

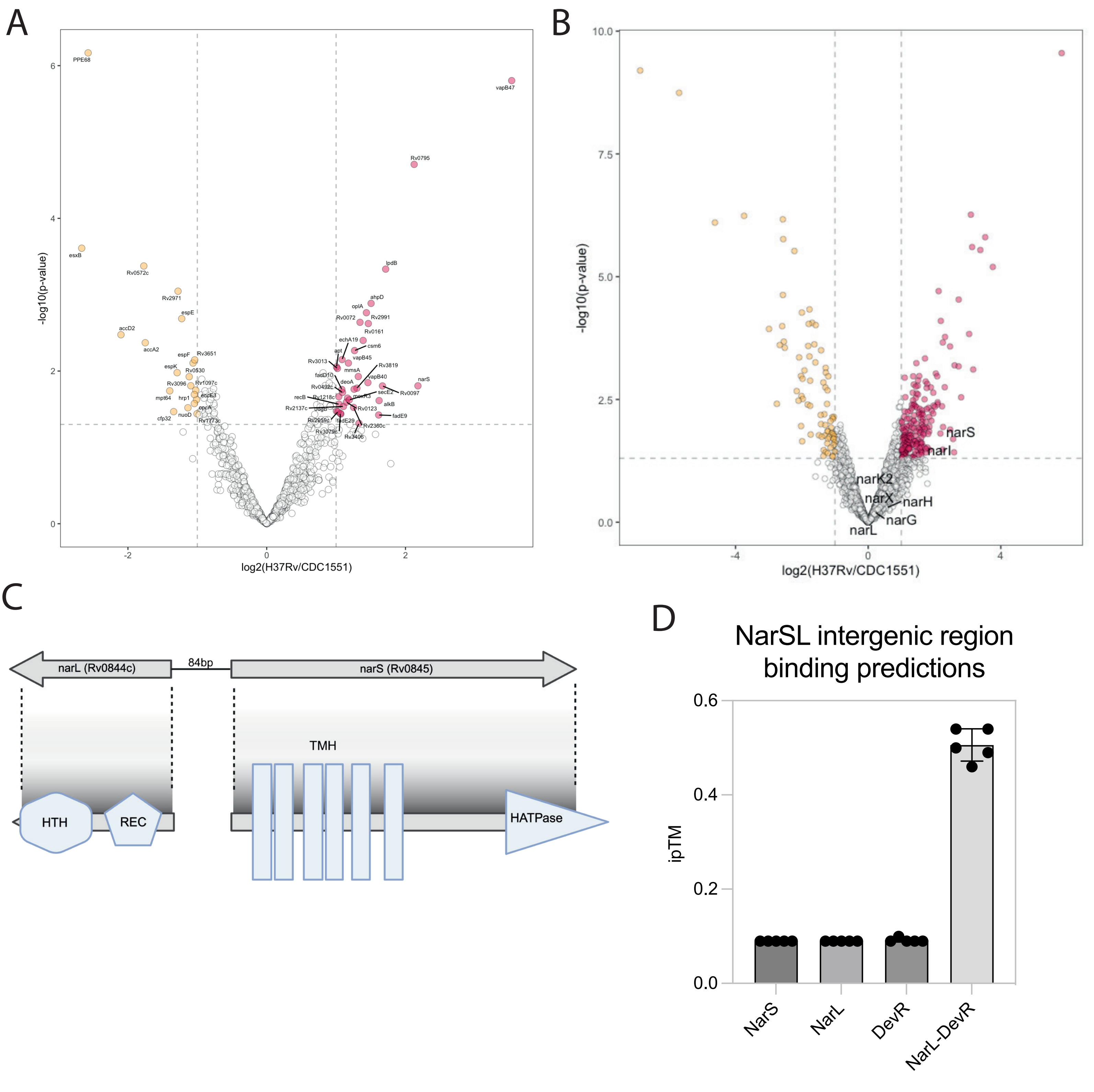

Supplementary Table 1

A

| Uniprot IDs | Gene name | Fasta headers | Average iBAQ |
|---|---|---|---|
| O05458 | mycP2 | Mycosin-2 | 0,650542948 |
| O07171 | Rv0121c | Conserved protein | 0,916153744 |
| O33347 | relF | Antitoxin RelF | -2,321928095 |
| O50440 | chp2 | Diacyltrehalose acyltransferase Chp2 | -2,821928095 |
| P71983 | Rv1725c | HTH hxlR-type domain-containing protein | -0,736965594 |
| P9WHI5 | recO | DNA repair protein RecO | -1,321928095 |
| P9WI49 | PPE1 | Uncharacterized PPE family protein PPE1 | -1,821928095 |
| P9WMD3 | Rv1353c | HTH-type transcriptional repressor Rv1353c | -0,881285397 |
| P9WPL5 | cyp142 | Steroid C26-monooxygenase | 1,428960383 |
| Q10884 | hycE | Possible formate hydrogenase HycE | 0,481737062 |

B

| Uniprot IDs | Gene name | Fasta headers | Average iBAQ |
|---|---|---|---|
| O53466 | Rv2020c | Type ISP restriction-modification enzyme LLaBIII C-terminal specificity domain-containing protein | 1,94734114 |
| O53837 | Rv0826 | ER-bound oxygenase mpaB/mpaB'/Rubber oxygenase catalytic domain-containing protein | 1,839458281 |
| P96391 | Rv0209 | Uncharacterized protein | -2,052809788 |
| P96937 | Rv0648 | Alpha-mannosidase | -1,598619761 |
| P9WH15 | rlmN | Probable dual-specificity RNA methyltransferase RlmN | 4,321560142 |
| P9WIF7 | PE_PGRS24 | Uncharacterized PE-PGRS family protein PE_PGRS24 | -2,029446845 |
| P9WJI5 | nat | Arylamine N-acetyltransferase | 0,555001678 |
| P9WL99 | Rv2561/Rv2562 | Uncharacterized protein Rv2561/Rv2562 | 1,450852289 |
| P9WMB7 | mftR | Putative mycofactocin biosynthesis transcriptional regulator MftR | -0,245655198 |
| Q50616 | Rv1817 | Possible flavoprotein | -1,486965594 |
| Q79FX5 | fabD2 | Possible malonyl CoA-acyl carrier protein transacylase FabD2 | 0,801271021 |

Supplementary Table 2

| Protein name | Fasta headers | Log2(Fold Change) | -LOG10(p-value) |
|---|---|---|---|
| vapB47 | Antitoxin VapB47 | 3.53042158876429 | 5.80402570041948 |
| narS | Sensor histidine kinase NarS | 2.17923147131284 | 1.80639434349621 |
| Rv0795 | Insertion element IS6110 uncharacterized 12.0 kDa protein | 2.12475260220535 | 4.70654818235583 |
| lpdB | Probable oxidoreductase | 1.71403369605134 | 3.33473643416101 |
| Rv0097 | (3R)-3-[(carboxymethyl)amino]fatty acid oxygenase/decarboxylase | 1.66829674776709 | 1.80505779922706 |
| alkB | Probable transmembrane alkane 1-monooxygenase AlkB | 1.61952629368701 | 1.61317787391054 |
| fadE9 | Probable acyl-CoA dehydrogenase FadE9 | 1.6133518948802 | 1.42436804612944 |
| ahpD | Alkyl hydroperoxide reductase AhpD | 1.50397540604292 | 2.88620538753985 |
| Rv2991 | Conserved protein | 1.46409525493654 | 2.62179158055122 |
| vapB40 | Antitoxin VapB40 | 1.45830078896233 | 1.84856776287058 |
| oplA | Probable 5-oxoprolinase OplA (5-oxo-L-prolinase) (Pyroglutamase) | 1.43656977222215 | 2.76390567452182 |
| Rv0161 | Possible oxidoreductase | 1.38923151791441 | 2.4006194445999 |
| Rv0072 | Uncharacterized ABC transporter permease Rv0072 | 1.34433589875872 | 2.6382477092602 |
| Rv3406 | Alpha-ketoglutarate-dependent sulfate ester dioxygenase | 1.32944420345229 | 1.31603238928549 |
| mmsA | methylmalonate-semialdehyde dehydrogenase (CoA acylating) | 1.31965372764566 | 1.92829587433021 |
| Rv3819 | Uncharacterized protein | 1.29995827174589 | 1.77567599514686 |
| csm6 | CRISPR system endoribonuclease Csm6 | 1.2659162255244 | 2.2678843311412 |
| deoA | Thymidine phosphorylase | 1.25932752458787 | 1.76166963049496 |
| Rv2360c | Uncharacterized protein | 1.25207984011411 | 1.52408280618547 |
| secE2 | Calcium dodecin | 1.19043846362848 | 1.62235376638861 |
| vapB45 | Putative antitoxin VapB45 | 1.17722502807895 | 2.10446677983589 |
| moxR3 | Probable methanol dehydrogenase transcriptional regulatory protein MoxR3 | 1.16501126426475 | 1.64539869780878 |
| Rv0123 | DNA-binding protein | 1.14811912116317 | 1.59303928294832 |
| Rv2137c | Uncharacterized protein | 1.10528582175778 | 1.53620574026899 |
| Rv0492c | Uncharacterized GMC-type oxidoreductase Rv0492c | 1.09212990475171 | 1.71955139893519 |
| echA19 | Enoyl-CoA hydratase EchA19 | 1.08588038092594 | 2.15017718034846 |
| fadD10 | Medium/long-chain-fatty-acid--[acyl-carrier-protein] ligase FadD10 | 1.08275344828441 | 1.75902569271951 |
| fadE29 | Acyl-CoA dehydrogenase FadE29 | 1.06831878480502 | 1.43791554851711 |
| Rv3079c | Conserved protein | 1.06205775145718 | 1.44474593163207 |
| recB | RecBCD enzyme subunit RecB | 1.04329218871908 | 1.56493207429172 |
| Rv1218c | Multidrug efflux system ATP-binding protein Rv1218c | 1.03852169111065 | 1.66432249468829 |
| Rv2959c | Rhamnosyl O-methyltransferase | 1.02629513281589 | 1.46796515846776 |
| Rv3013 | Conserved protein | 1.01921601796654 | 2.03401004755092 |
| apt | Adenine phosphoribosyltransferase | 1.00624434873909 | 2.05362948984038 |
| udgB | Type-5 uracil-DNA glycosylase | 1.00367182709413 | 1.47678803094239 |

### Supplementary Table 3

| Protein name | Fasta headers | Log2(Fold Change) | -LOG10(p-value) |
|---|---|---|---|
| esxB | ESAT-6-like protein EsxB | -2.66614398487351 | 3.6091062880844 |
| PPE68 | PPE family immunomodulator PPE68 | -2.5738292526439 | 6.16738091675419 |
| accD2 | Probable biotin-dependent acyl-coenzyme A carboxylase beta2 subunit | -2.10061619629601 | 2.47548130110894 |
| Rv0572c | Uncharacterized protein Rv0572c | -1.77152119916825 | 3.37590155912867 |
| accA2 | Probable acetyl-/propionyl-coenzyme A carboxylase alpha chain (Alpha subunit) AccA2 | -1.75046363117555 | 2.37068746689103 |
| mpt64 | Immunogenic protein MPT64 | -1.39756055803398 | 1.74060197064803 |
| cfp32 | Putative glyoxylase CFP32 | -1.34111150011474 | 1.46817231355069 |
| espK | ESX-1 secretion-associated protein EspK | -1.29091403350802 | 1.97864677142218 |
| Rv2971 | Aldo-keto reductase Rv2971 | -1.27816642361563 | 3.04594412386888 |
| espE | ESX-1 secretion-associated protein EspE | -1.22517118875644 | 2.6865234911556 |
| nuoD | NADH-quinone oxidoreductase subunit | -1.13551337254187 | 1.51907679138612 |
| Rv0530 | Conserved protein | -1.11707866108325 | 1.92613498170886 |
| Rv3096 | 1,4-beta-xylanase | -1.0939854534078 | 1.80745700119004 |
| espF | ESX-1 secretion-associated protein EspF | -1.06192109023663 | 2.10515558727501 |
| opcA | OXPP cycle protein OpcA | -1.04278405226569 | 1.57443580001827 |
| hrp1 | Hypoxic response protein 1 | -1.04261835503554 | 1.69269000335838 |
| Rv3651 | Rv3651-like N-terminal domain-containing protein | -1.03770557238453 | 2.14607214214457 |
| Rv1097c | Probable membrane glycine and proline rich protein | -1.02313182410315 | 1.74964750501892 |
| eccE3 | ESX-3 secretion system protein EccE3 | -1.01168924499053 | 1.62520207013748 |
| Rv1773c | Probable transcriptional regulatory protein | -1.00446865523486 | 1.44015301480426 |

Supplementary Figure 7

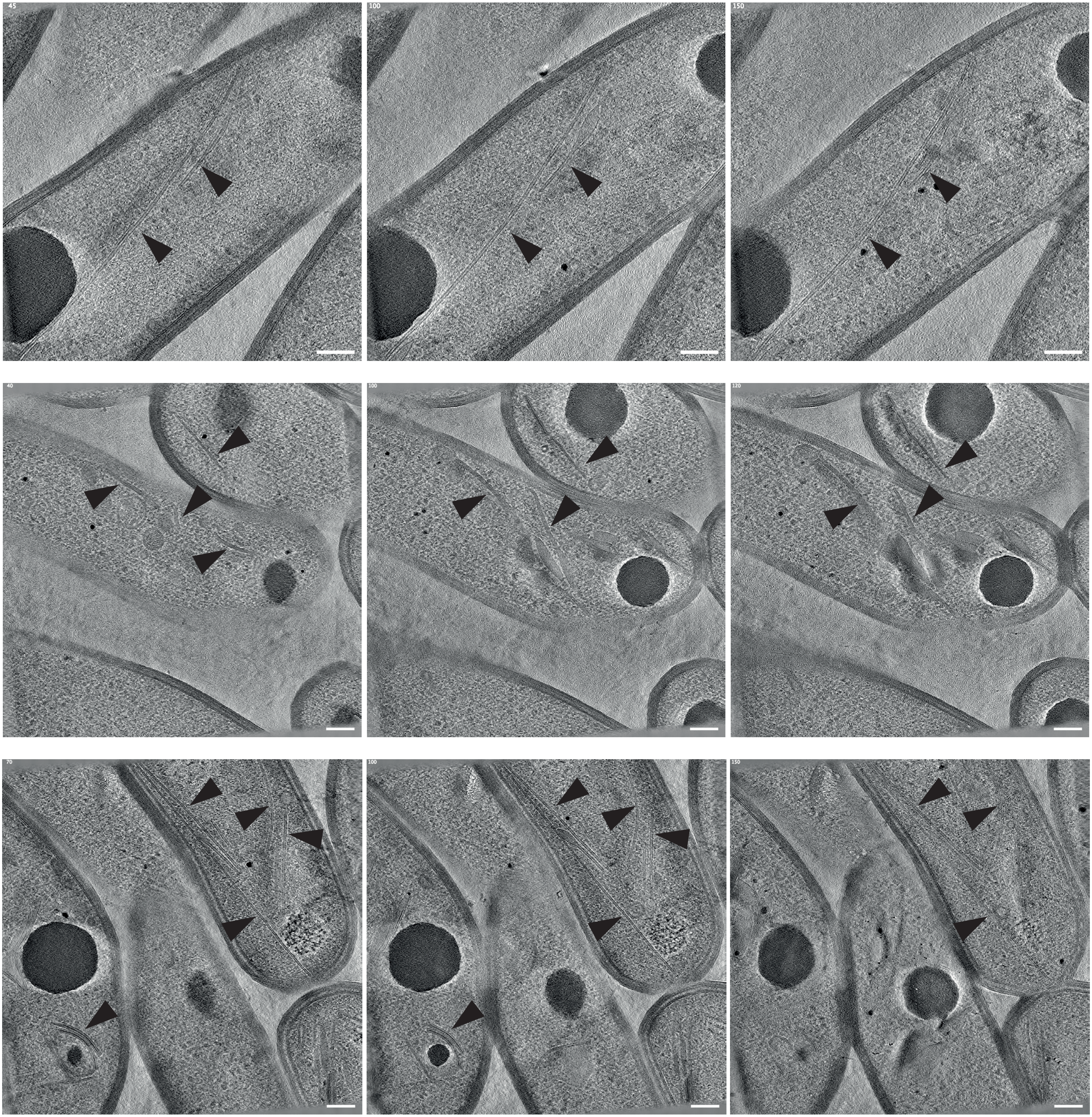

### Supplementary Figure 8

A

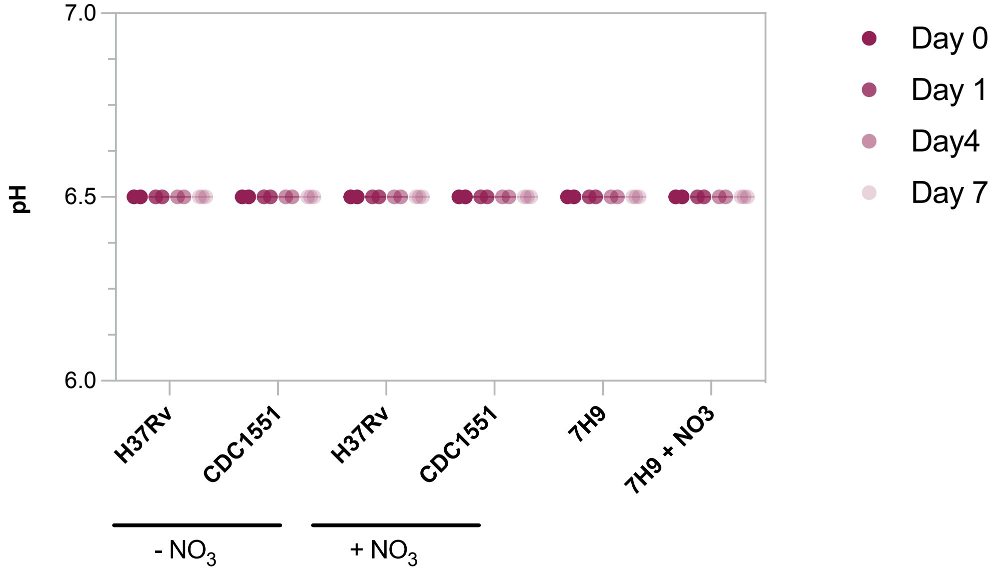

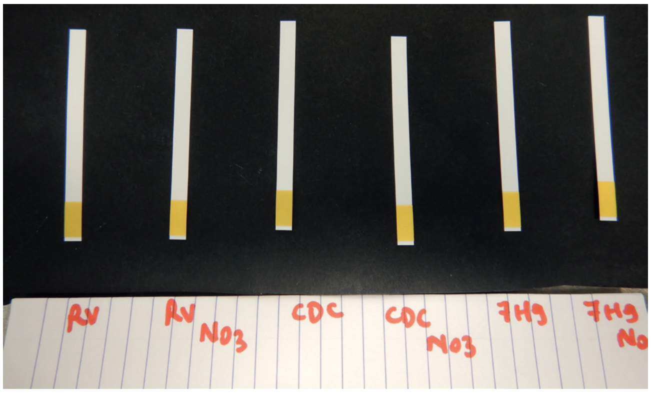

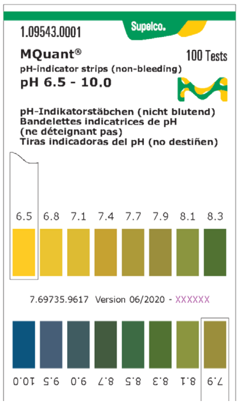

B

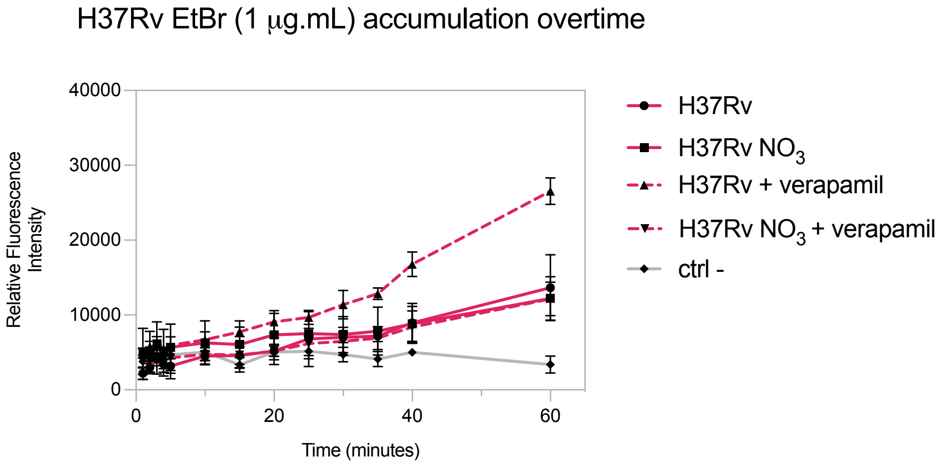

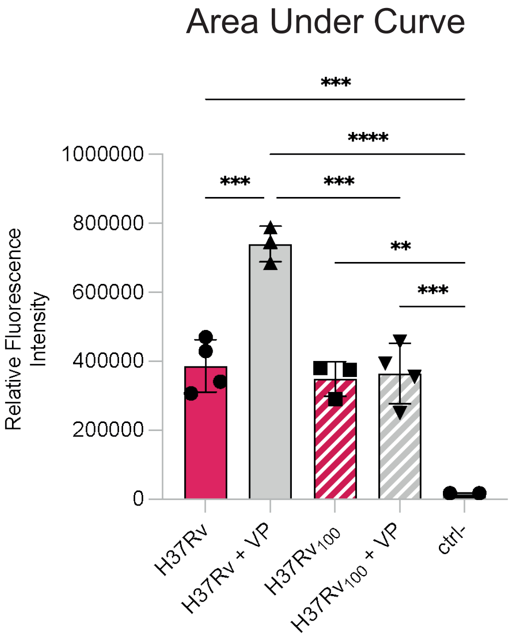

### Supplementary Figure 9

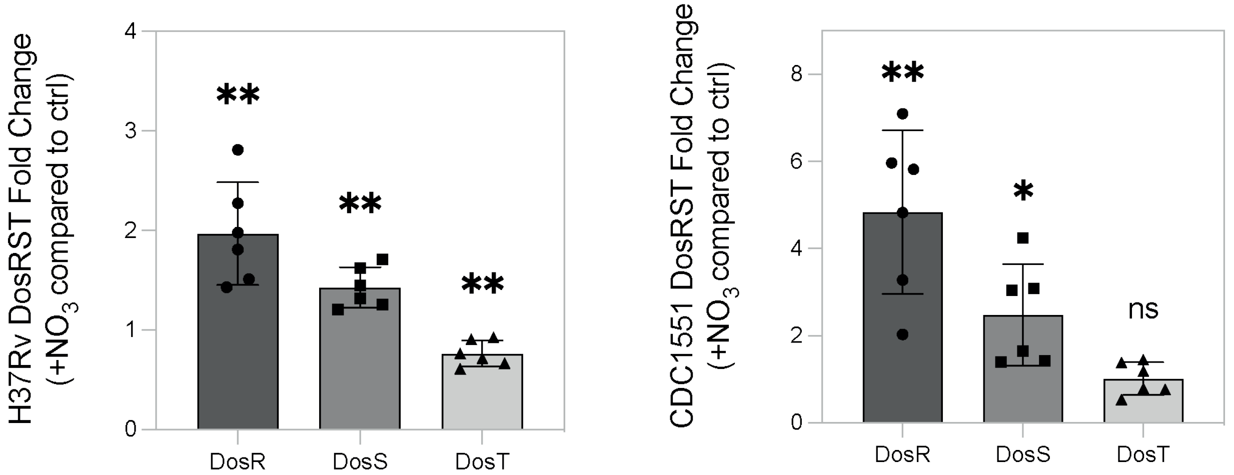

Supplementary Figure 10

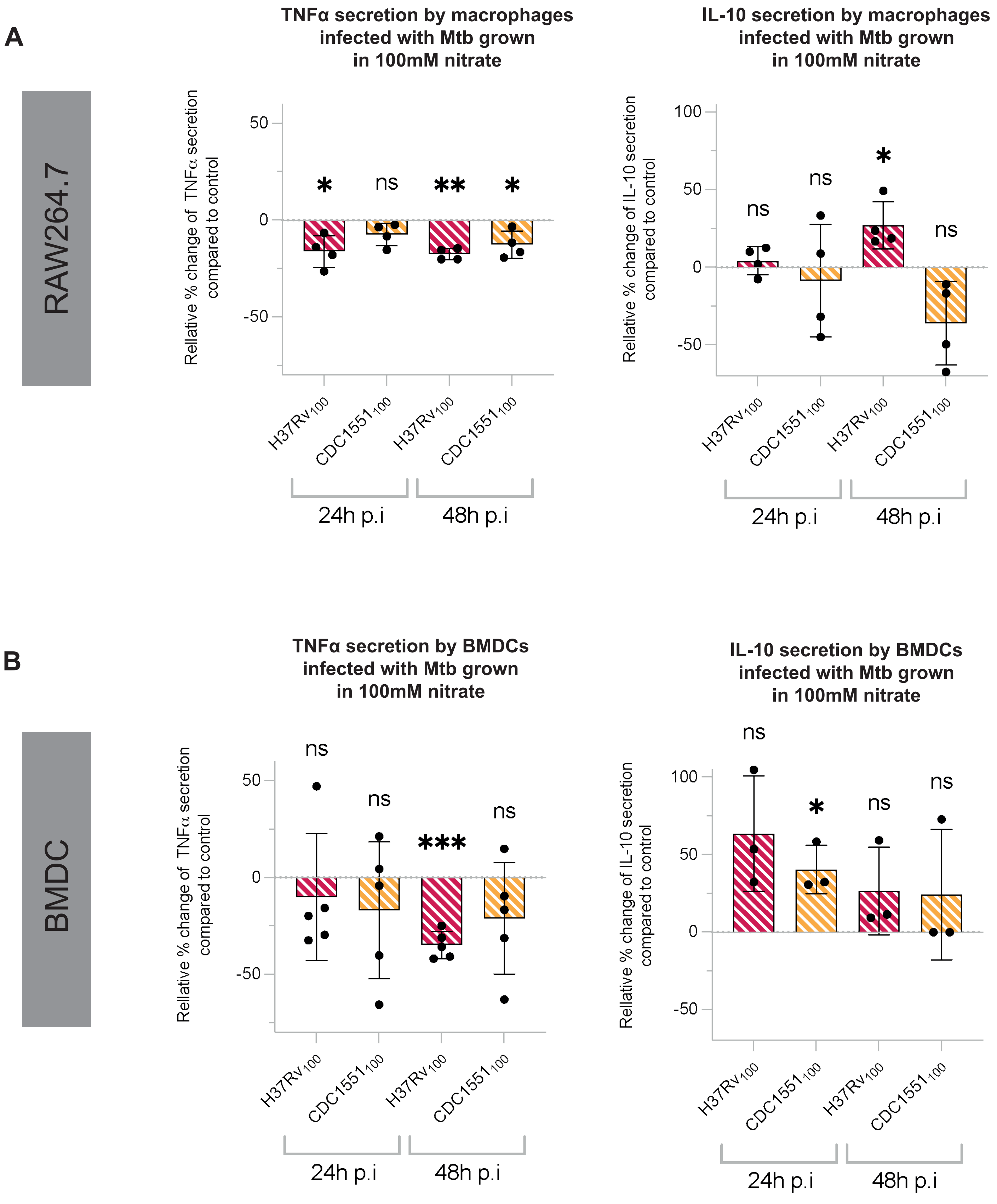

Supplementary Table 4

| Primer | Sequence | Purpose |
|---|---|---|
| NarS_R | gtttgggcggtcgccatgagta | qPCR NarS (Rv0845) |
| NarS_F | tggtagttccacgcagacatgggtc | |
| NarL_R | acgccgagcttctcgtacaaccg | qPCR NarL (Rv0844c) |
| NarL_F | tgcgcgtggtggtcggcga | |
| DosS_R | gacaaagtgcaatacccgatgc | qPCR DosS (Rv3132c) |
| DosS_F | tacgcctgcacgagctgct | |
| DosR_R | caactccattcccttgatgtctttga | qPCR DosR (Rv3133c) |
| DosR_F | tcttcttggtcgatgaccacgaggt | |
| DosT_R | cacctcgtcgtcatcgctgaa | qPCR DosT (Rv2027c) |
| DosT_F | attcgtctacgaggggatcgacga | |
| ahpD_R | tgtggccagcgcttgtgcaac | qPCR ahpD (Rv2429) |
| ahpD_F | ctccccgagtacgccaaagacatca | |
| Rv3093c_R | agcgctcgcagagccaccc | qPCR Rv3093c |
| Rv3093c_F | taccgttctggcttgaccgcc | |
| egtB_R | agatctggcggcgatacgggtg | qPCR egtB (Rv3703c) |
| egtB_F | gagcagctggcttgtcatctggc | |
| Rv0161_R | gtccaacgccgccttgac | qPCR Rv0161 |
| Rv0161_F | accacaccggccgctatc | |
